# Proteome-wide identification and modeling of interactions between transactivation domains and arginine-glycine-rich regions

**DOI:** 10.64898/2026.04.13.717676

**Authors:** Yukti Khanna, Arijana Hajdarević, Johanna Pirchner, Sinem Usluer, Anastasia Rakhimbekova, Iva Pritišanac, Sören von Bülow, Kresten Lindorff-Larsen, Tobias Madl

## Abstract

Transcription factors (TFs) and RNA-binding proteins (RBPs) coordinate gene expression across transcriptional and post-transcriptional layers, yet the principles that govern their direct physical coupling, especially through intrinsically disordered regions, remain unclear. Here we combine proteome-scale interaction mapping, disordered-region annotation, coarse-grained simulations and sequence-based prediction to dissect a prevalent TF-RBP interface mediated by acidic/hydrophobic transactivation domains (TADs) and arginine-glycine-rich (RG/RGG) regions. Network analysis reveals a global enrichment of RBP partners among TF interactions and identifies TF and RBP hubs that bridge transcriptional regulation with RNA-centered pathways. Using a sequence grammar enriched in acidic and aromatic residues, we define 230 RG/RGG-binding TAD-like segments across 190 TFs and we map 1,008 compact RG/RGG regions across 823 RBPs based on proteome-wide motif spacing. Coarse-grained simulations (CALVADOS) of representative TAD-RGG pairs quantify interaction propensities and indicate that association is primarily driven by electrostatic complementarity and charge patterning, with sequence “stickiness” modulating interaction strength. Using a hybrid machine-learning model we predicted simulated interaction strengths from a compact, interpretable set of features and extrapolate these rules to the full combinatorial space, enabling systematic prioritization of candidate TF-RBP couplings. To validate these predictions experimentally, we used NMR titration experiments on a subset of TAD-RGG pairs spanning the predicted affinity range, which showed agreement between predicted affinities and NMR-derived dissociation constants. Together, our results support a predominantly electrostatic mode of association and establish a quantitative framework for identifying and prioritising TF-RBP partnerships, revealing how complementary sequence grammars within disordered regions couple transcriptional regulation to RNA processing and transport.

## Introduction

Intrinsically disordered regions (IDRs) are widespread and functionally indispensable components of the eukaryotic proteome. Unlike structured domains that adopt a stable tertiary conformation, IDRs populate dynamic conformational ensembles, whose properties are determined by both sequence composition and patterning [1]. These regions are particularly enriched in regulatory proteins such as transcription factors (TFs) and RNA-binding proteins (RBPs), where they facilitate context-dependent interactions that modulate gene expression. These interactions are mediated by short linear motifs, post-translational modifications, and distributed interaction-prone residues (“stickers”) along the disordered chain [1,2].

Crucially, IDRs are not compositionally random. Proteome-scale analyses reveal that disordered sequences can be categorized into distinct “molecular grammar” classes, defined by specific amino acid patterning and spacing rules [3]. Complementary approaches using protein language models and evolutionary conservation further highlight sequence-encoded motifs associated with phase separation and condensate formation [4]. Beyond motif identification, machine learning is increasingly employed to generate conformational ensembles of IDRs, offering a predictive framework that links sequence features to structural and interaction propensities [5]. These findings collectively support the view that IDR behavior can, to a large extent, be inferred from primary sequence.

In contrast to folded domains, IDRs often engage in multivalent, “fuzzy” interactions driven by weak forces, including electrostatic interactions, π–π and cation–π contacts, and transient hydrophobic associations [1,6]. These same interaction modalities also underpin the formation of biomolecular condensates, such as those formed by prion-like domains of RBPs and by transcriptional coactivators [7–9]. Nevertheless, large-scale interactome datasets indicate that IDR-mediated contacts can appear promiscuous, raising a fundamental challenge: distinguishing biologically meaningful interaction grammars from generic “stickiness” at the proteome scale. Recent efforts to predict IDR–IDR interactions directly begin to address this challenge, though generalizable and quantitative pairing rules remain elusive [10].

To rationalize disordered behavior and interaction specificity, several complementary descriptor systems have been proposed. Polymer-physics metrics, such as net charge per residue (NCPR) and the fraction of charged residues, govern chain compaction and responsiveness to ionic conditions. Charge patterning introduces another layer of control: sequences with segregated patches of opposite charge tend to be more compact, whereas sequences with more uniformly distributed charges are typically more expanded, depending on net charge and solution conditions [11,12]. Tools such as CIDER and localCIDER enable systematic mapping of these sequence-to-ensemble relationships [12], while grammar-based resources such as GIN (Grammars Inferred using NARDINI+) offer proteome-wide classification schemes that incorporate composition and length [3,13]. Simultaneously, IDRs can act as molecular tethers and sensors of cellular chemistry, responding to physicochemical changes in their environment via modulations in conformational ensembles [14]. Together, these perspectives promote a sequence-level understanding of how TF and RBP IDRs engage in selective, multivalent interactions, rather than indiscriminate binding.

Two disordered module types are especially well positioned to engage via physicochemical complementarity: acidic/hydrophobic transactivation domains (TADs) in TFs and arginine-glycine-rich (RG/RGG) regions in RBPs. These modules combine pronounced charge asymmetry with low-complexity sequence composition, suggesting that their interactions may be both selective and tunable by sequence context. Here, we test this hypothesis by systematically identifying candidate TADs and compact RG/RGG regions and quantifying their interaction propensities using coarse-grained simulations together with sequence-based predictive modeling.

TADs are short (∼10–50 residues), compositionally diverse IDRs that recruit coactivators and chromatin-modifying complexes to drive transcription [4]. Although historically classified by dominant residue type, most functional TADs are better characterized as acidic/hydrophobic hybrids, featuring interspersed bulky hydrophobic or aromatic residues and acidic residues that enhance solubility and enable binding [15]. TADs typically remain disordered upon binding, forming fuzzy interfaces and transient secondary structure, with interaction mechanisms shaped by sequence-encoded conformational propensities and local context [16–19]. Some TADs also function as conformational switches that regulate coactivator engagement [20–22]. In addition, negatively charged residues can suppress degradation mediated by exposed hydrophobic patches, helping explain the recurrent coupling of acidity with hydrophobic motifs in activation domains [23,24]. While many TADs have been associated with the formation of coactivator-rich condensates [8,9], phase separation does not necessarily equate to functional transcriptional output, as condensate formation and activity can be partially uncoupled [25].

Conversely, many RBPs contain disordered RGG/RG regions (“RGG boxes”) that are rich in arginine and generally positively charged [26–28]. These regions support weak, multivalent interactions with diverse partners, including RNA, folded domains, and other IDRs, through a combination of electrostatic and π-mediated contacts [27,28]. RG-rich segments are frequently implicated in condensate-linked assemblies such as stress granules and other membrane-less compartments [29–32], and their interaction propensities can be tuned by post-translational modifications, particularly arginine methylation [33–36]. Despite their conserved motifs, RGG regions exhibit wide variability in architecture, differing in motif density, spacing, and flanking residue composition, leaving open the question of how region architecture and motif organisation shape partner specificity and interaction strength [27,28].

Several case studies illustrate direct TF–RBP interactions mediated by disordered regions. For example, in the DNA damage response, the phosphorylated TAD2 domain of p53 binds the RG-rich GAR domain of MRE11 [37]. The RBP ALYREF can function as a TF coactivator in multiple contexts, including through an ALYREF–MYCN complex and through modulation of LEF-1/AML-1 and E2F2 activity [38–40]. FUS, a well-studied RBP, has also been implicated in transcriptional regulation via its involvement in coactivator-enriched nuclear assemblies [41]. However, these observations remain difficult to generalize, and a quantitative framework linking disordered-region sequence features to TF–RBP interaction propensity at scale is still missing.

To address these gaps, we integrate proteome-scale interaction data, coarse-grained simulations, and sequence-based modeling. First, we construct a TF-centered interaction network to quantify TF–RBP connectivity patterns. Second, we use CALVADOS v2 simulations to estimate interaction strengths for representative TAD– RGG pairs [42].

More broadly, our results support a rule-based view of disorder-mediated interactions in which sequence-encoded physicochemical complementarity can confer selective and tunable binding. By enabling quantitative ranking of candidate TF–RBP pairs from sequence, this framework provides an actionable guide for experimental follow-up, highlighting which partners to test first and which to prioritize next, and offers a general template to map interaction propensities between other protein groups.

## Results

### Transcription factors preferentially interact with RNA-binding proteins

To quantitatively assess the extent to which transcription factors [43] engage RNA-binding proteins [44] within the human protein–protein interaction (PPI) network, we analyzed 10,043 unique TF-centered interactions derived from BioGRID and I2D [45], encompassing 1,464 curated TFs. TFs exhibited significantly higher fractions of RBP partners than any other protein class (Wilcoxon rank-sum tests; Benjamini–Hochberg q ≤ 10^− 2^□; Fig. 1A; Supplementary Table S1). A complete per-TF breakdown of partner composition and RBP interactor lists is provided in Supplementary Table S1 (sheets S1A–S1B). To further evaluate this enrichment of TF–RBP interactions, we classified interaction partners of each TF into four functional categories: RBPs, other TFs, protein-modifying enzymes (PMEs), and randomly sampled proteins. Because the RBP, TF, and PME reference sets differ in size (with RBPs being the largest), we assessed enrichment as a per-TF partner fraction and used a random protein set size-matched to the RBP set as a conservative baseline. RBPs consistently accounted for the largest proportion of partners per TF, with a median near 25%, while TFs, PMEs, and random proteins each contributed substantially smaller fractions (Fig. 1A). The distribution of RBP interaction frequencies was significantly elevated compared to all other categories (Mann–Whitney U/Wilcoxon rank-sum tests; all BH-corrected q ≤ 10^− 2^□; Fig. 1A; Supplementary Table S1). These findings indicate that TFs are not uniformly connected and show a pronounced enrichment of RBP partners in the curated PPI dataset.

**Figure 1.**
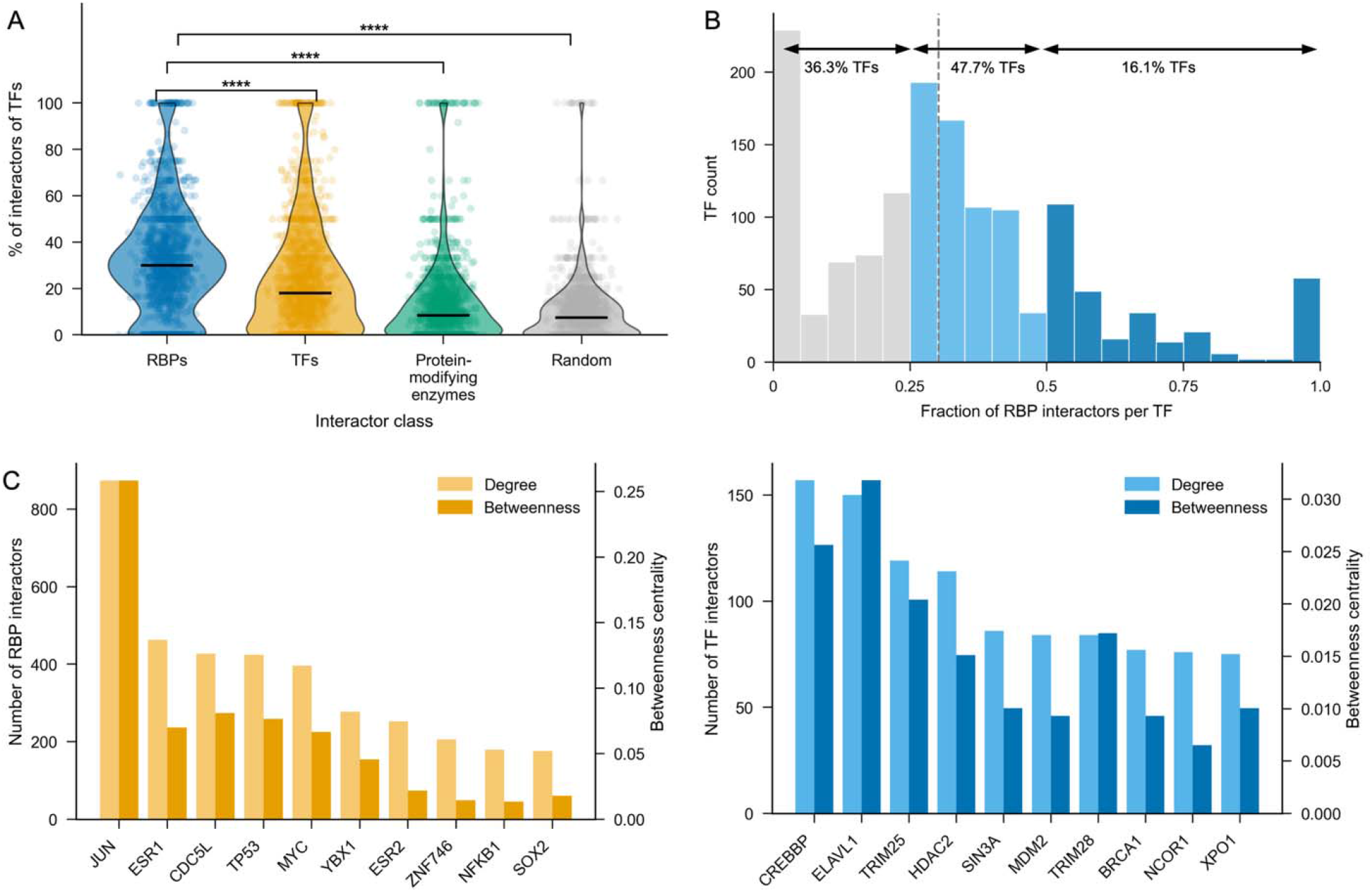
Transcription factors preferentially interact with RNA-binding proteins. (A) Violin plots showing the per-transcription factor percentage of interaction partners classified as RBPs, other TFs, PMEs, or randomly sampled proteins. Each violin represents the distribution across 1,464 TFs, with the solid line indicating the proteome-wide frequency of each class. TFs exhibit a significantly higher fraction of RBP interactors compared to all other classes (Mann-Whitney U, all p ≤ 10^− 2^□). (B) Distribution of RBP interaction bias across the TF repertoire. The donut chart depicts the proportion of TFs for which <25%, 25–50%, or >50% of their interaction partners are RBPs. Approximately 16.1% of TFs are highly RBP-enriched, with more than half of their interactors classified as RBPs. (C) Top ten TF hubs ranked by betweenness centrality (dark bars, left axis), with corresponding numbers of RBP interactors shown (light bars, right axis). Canonical transcriptional regulators such as JUN, TP53, ESR1, and MYC are prominent network hubs with extensive RBP connectivity. (D) Top ten RBP hubs ranked by betweenness centrality (dark bars, left axis), with corresponding numbers of TF interactors shown (light bars, right axis). Key proteins annotated as RBPs in EuRBPDB, including ELAVL1, CREBBP, TRIM25, and HDAC2 exhibit both high centrality and broad TF connectivity. Together, panels C and D identify a cohort of multifunctional hub proteins that mediate connectivity between transcriptional and RNA-regulatory subnetworks within the cuarted interactome.

This interaction bias was observed to be widespread but heterogeneously distributed across the TF repertoire (Fig. 1B). Specifically, 36.3% of TFs had less than 25% of their partners classified as RBPs, 47.7% displayed an intermediate interaction level (25– 50%), and 16.1% were characterized by a predominant RBP interaction profile, with over half of their interactors being RBPs. These data suggest that RBPs represent a major, and in a subset of TFs, the principal class of interaction partners. Our network-level enrichment analyses were performed on curated PPIs, and RBP assignment was based on EuRBPDB; because these data do not employ degree-preserving null models or modality-matched controls and may include multifunctional regulators, the observed enrichment should be interpreted as a robust trend in available curated datasets rather than a degree-controlled estimate of class-specific binding preference.

To determine whether this enrichment is concentrated within specific regions of the network, we computed betweenness centrality for each TF and RBP and examined their degree of cross-class connectivity (Fig. 1C, D). Prominent transcriptional regulators such as TP53, JUN, ESR1, and MYC exhibited high betweenness centrality along with numerous RBP interactors, indicating their role as TF hubs. Conversely, proteins annotated as RBPs in EuRBPDB, including ELAVL1, CREBBP, HDAC2, and TRIM25 demonstrated reciprocal patterns of high centrality and extensive TF connectivity; this class should be interpreted as an annotation-defined RBP set that includes canonical and non-canonical RNA-binding proteins. These findings identify a defined subset of proteins as prominent network bridges linking transcriptional regulation with RNA processing and signaling subnetworks within the curated interactome. Centrality metrics for all TF and RBP nodes are summarized in Supplementary Table S1 (sheets S1C– S1D).

In summary, transcription factors are enriched for RBP partners in curated PPIs; enrichment is robust to within-TF fractioning but requires degree-controlled nulls for causal inference. This connectivity is not uniformly distributed but is concentrated in a cohort of central hubs that connect DNA- and RNA-associated regulatory processes.

### Sequence composition and patterning distinguish R-TADs and define compact RG regions for partner selection

Having established that transcription factors show a strong preference for interacting with RNA-binding proteins in the human protein–protein interaction network, we next asked which parts of TF sequences could provide the physical interface for these contacts (Fig. 1; Supplementary Table S1). Many RBPs recognise partners via basic arginine-glycine–rich regions, so we hypothesised that complementary, negatively charged activation domains might act as cognate binding modules. As a minimal model for RGG/RG regions, we used a synthetic [RRGG]_7_ peptide, which recapitulates the high density and spacing of RG motifs typical of RBP low-complexity domains (Fig. 2A). Direct binding assays with this peptide identified four acidic activation domains, LEF1 TAD, FOXM1 TAD, the N-terminal TAD of p53 and the FOXO4 TAD (also named conserved region 3/CR3), as robust interactors, which we term “seed R-TADs” (RG/RGG-binding transcriptional activation domains) (Fig S1A-D). Because acidic activation domains are often conserved within TF families, we expanded this seed set by including homologous acidic segments from related FOX/TCF family members (TCF4, FOXP2, FOXO1, FOXO3 and FOXO6), yielding a focused panel of TFs whose activation domains can recognise RG/RGG-like peptides (Supplementary Table S2, sheet S1A).

**Figure 2.**
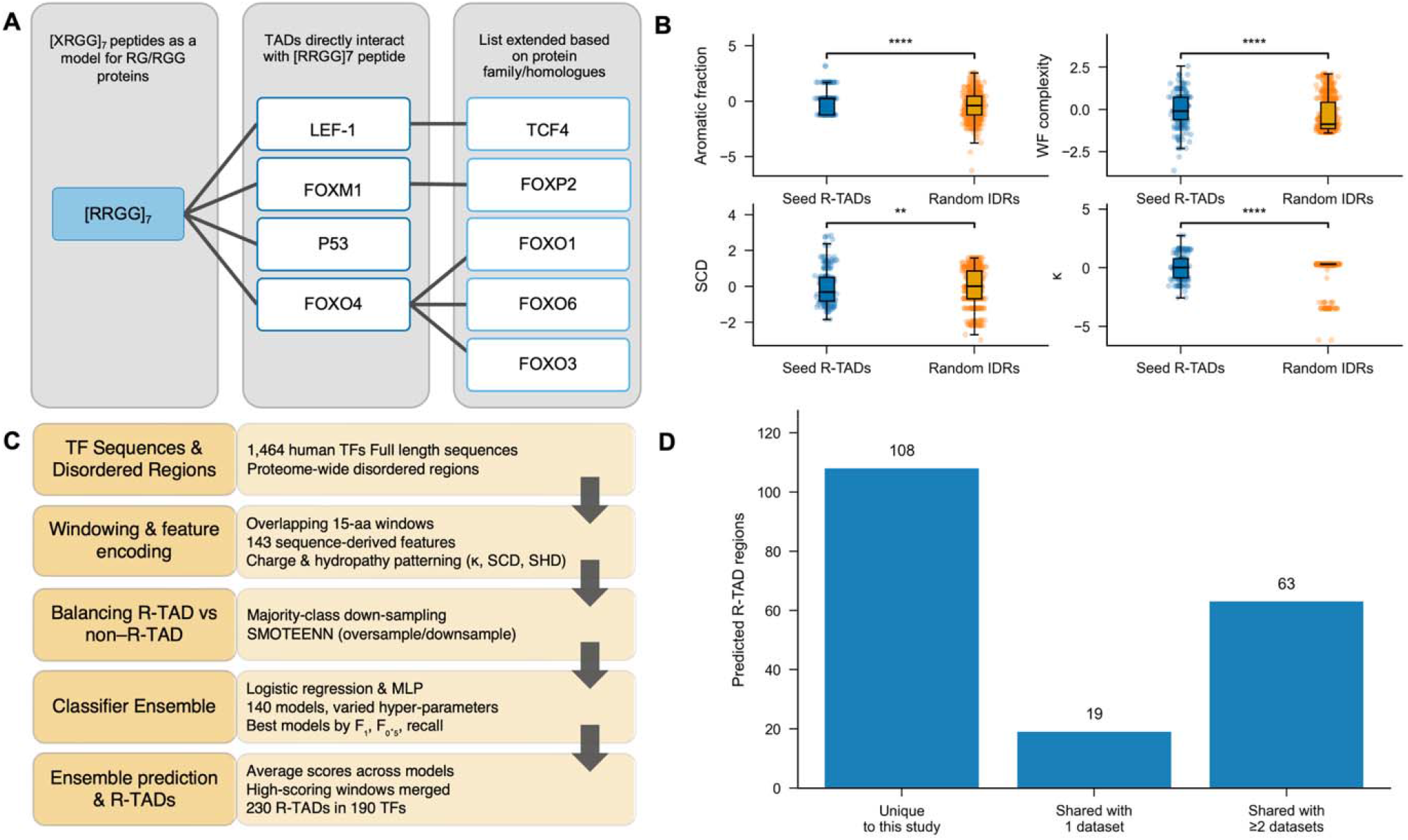
Acidic and hydrophobic sequence features define transactivation domains. (A) Selection of seed R-TADs using an RG/RGG model peptide. Schematic of the strategy used to define RG/RGG-binding activation domains. A synthetic [XRGG]_7_ peptide (illustrated as [RRGG]_7_) serves as a generic model for RG/RGG regions in many RBPs. Binding assays identified four acidic activation domains, LEF1 TAD, FOXM1 TAD, p53 TAD and the FOXO4 TAD as robust interactors (“seed R-TADs”). Homologous acidic regions from related FOX/TCF family members (TCF4, FOXP2, FOXO1, FOXO3 and FOXO6) were added to generate an extended panel of RG/RGG-binding activation domains. (B) Sequence features of seed R-TADs versus random TF IDRs. Box-and-whisker plots comparing feature values between windows from seed R-TADs (blue) and length-matched disordered windows from other TF regions (orange). Shown are aromatic content (fraction of Phe/Tyr/Trp), aromatic patterning (WF_complexity, a measure of local sequence repetitiveness; lower values indicate more repetitive/clustered aromatics), and two charge-patterning metrics (SCD and κ; lower values indicate more mixed charge distributions). Points are individual windows; boxes show medians and interquartile ranges; whiskers extend to 1.5× IQR. Seed R-TADs are enriched in aromatics, have lower WF_complexity and display more mixed charge patterning than random TF IDRs (Wilcoxon tests; *** or **** indicate significant differences). (C) Machine-learning workflow for proteome-wide identification of R-TADs. Schematic of the classification pipeline. TF sequences were segmented into overlapping 15-residue windows and encoded by 143 sequence-derived features [48], including composition, charge, hydropathy and patterning descriptors (κ, SCD, SHD). An ensemble of logistic-regression and MLP classifiers was trained using different hyper-parameter and resampling schemes. Class imbalance between TAD-like and non-TAD windows was addressed by combining majority down-sampling with SMOTEENN. Predictions from the best-performing models were averaged, and overlapping high-scoring windows were merged to define R-TAD segments. (D) Support for predicted R-TAD regions across published AD/TAD datasets. Predicted R-TAD regions were grouped according to the number of external AD/TAD datasets (Soto 2022 [50], Kotha & Staller 2023 high-confidence set, Kotha & Staller 2023 GSL library [15], Staller 2022 [47]) in which a corresponding mapped region was detected: “Unique to this study” indicates that no corresponding mapped region was detected in any prior dataset; “Shared with 1 dataset” indicates that a corresponding mapped region was detected in exactly one prior dataset; and “Shared with ≥2 datasets” indicates that corresponding mapped regions were detected in two or more prior datasets.

To test whether these RG/RGG-binding transactivation domains (R-TADs) share a recognizable sequence grammar, we compared their composition and patterning to length-matched, disordered regions drawn at random from other TFs. Each region was encoded by sequence features capturing aromatic content, charge distribution and local hydropathy (Methods; underlying values and statistics in Supplementary Table S2, sheets S1C–S1D). As a seed-level comparison, length-matched TF IDRs provide a conservative baseline to highlight qualitative differences in aromatic content and charge mixing. Seed R-TADs displayed a coherent, distinctive profile relative to random TF IDRs (Fig. 2B): they were enriched in aromatic residues (Phe/Tyr/Trp), consistent with potential cation–π contributions to binding Arg-rich partners, and these aromatics occurred in more repetitive, clustered patterns reflected by lower Wootton–Federhen (WF) sequence complexity, a metric that quantifies local sequence repetitiveness, with lower values indicating lower compositional complexity and greater clustering of similar residues. Charge-patterning metrics were shifted towards more mixed distributions of oppositely charged residues, indicating trends consistent with dispersed acidic residues rather than strongly segregated charge blocks.

We then generalised this sequence grammar to identify R-TADs proteome-wide. Using an ensemble classifier trained on 15-residue windows from disordered regions of 1,464 human TFs and encoded by 143 sequence-derived features (Fig. 2C; Methods), we identified 230 predicted R-TADs across 190 TFs. Overlapping high-scoring windows were merged into contiguous segments, and TFs carrying predicted R-TADs were compared with the full TF set in downstream analyses. (Supplementary Table S2, sheets S1B and S1E).

To benchmark this catalogue against previous efforts to map acidic/hydrophobic activation domains, we compared our predicted R-TAD sequences to four AD/TAD datasets derived from high-throughput functional assays (Soto 2022 [50]; Kotha & Staller 2023 high-confidence set; Kotha & Staller 2023 GSL library [15]; Staller 2022 [51]). Sequence-based mapping showed substantial but incomplete agreement across datasets: many predicted R-TADs aligned to previously reported activation-domain regions, whereas others lacked corresponding mapped segments and remained unique to our analysis (Fig. 2D, S2E; membership matrix in Supplementary Table S3, sheet S1A). Sequence-based mapping yield and quality across datasets are summarised in Supplementary Fig. S2A–B, with the underlying alignments provided in Supplementary Table S3 (sheets S3C–S3G; summary in sheet S3B). Several mapped activation-domain regions were supported across multiple datasets, whereas others were recovered in only a subset of studies. Notably, 108 predicted R-TADs regions in TFs were unique to our analysis, indicating that an RG/RGG-binding–oriented definition of activation domains captures both established and previously unannotated candidates.

Finally, we asked whether the broader set of predicted R-TADs preserves the compositional and patterning features observed for the experimentally characterised seeds. When we compared all predicted R-TAD segments to random disordered regions from other TFs, we again observed pronounced enrichment in aromatic content, lower WF complexity (more repetitive/aromatic clustering), and more mixed charge distributions (lower κ and a shifted SCD distribution under our implementation) in the R-TAD set (Supplementary Fig. S2C–D; underlying feature summaries in Supplementary Table S2, sheets S2C–S2D). The separation from random IDRs was at least as strong as for the seed panel, consistent with the classifier selecting windows that conform to this RG/RGG-binding grammar. Together, these analyses identify a compact class of acidic, aromatic, charge-dispersed activation domains, R-TADs, whose sequence features are consistent with providing negatively charged, sticker-rich interaction surfaces for the basic RG/RGG regions of RBPs highlighted by our network analysis.

To support downstream TF–RBP pairing analyses, we next established a consistent, rule-based definition of Arg–Gly–rich partner segments by identifying compact RG regions within disordered proteins. Although arginine and glycine are individually common in intrinsically disordered regions (IDRs), RG dipeptides are relatively rare under a random-composition model. Consistent with this expectation, background distributions across all disordered regions showed low RG densities, whereas curated RG/RGG proteins were strongly enriched for RG dipeptides per 100 residues (Fig. 3A; summary distributions in Supplementary Fig. S3A, most regions contain 10-20 RG repeats; curated RG/RGG disordered-region sequences and per-sequence metrics in Supplementary Table S4, sheet S4D).

**Figure 3.**
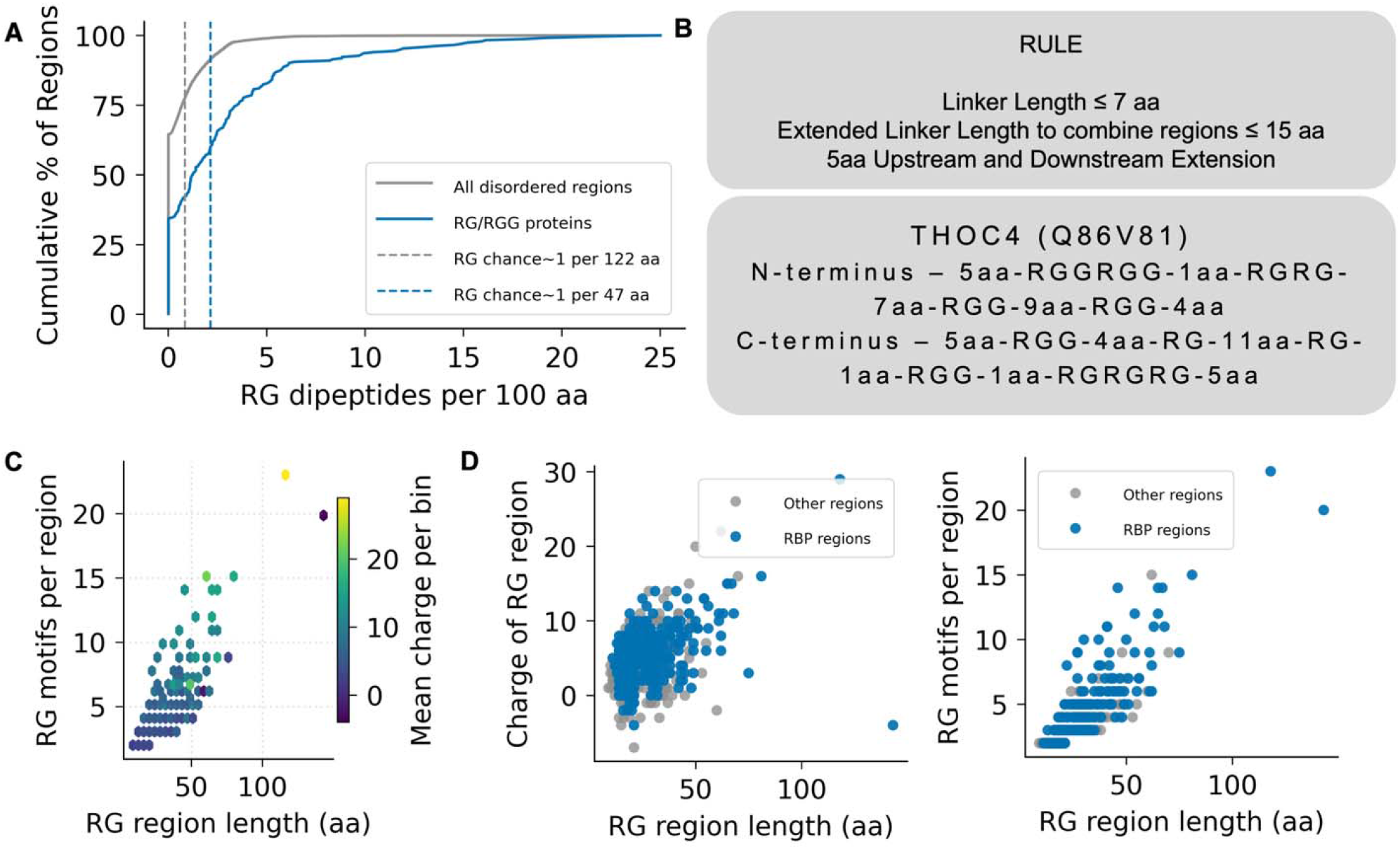
Rule-based definition of compact RG-rich regions and their proteome-wide properties. (A) Cumulative distributions of RG dipeptide density across disordered regions. The fraction of regions containing a given number of RG dipeptides per 100 amino acids is shown for all disordered regions in the proteome (grey) and for curated RG/RGG-containing proteins (blue). Dashed lines indicate the expected RG frequency under the corresponding background composition, expressed as approximately one RG dipeptide per indicated number of residues. (B) Top: Operational rule used to define compact RG regions. Consecutive RG motifs are grouped if separated by linkers of ≤7 residues; linkers up to 15 residues are permitted only to bridge two otherwise compact tracts. Each resulting region is extended by ±5 residues to preserve local sequence context [49]. Bottom: segmentation example from THOC4 (ALYREF), with RG-rich regions identified in both terminal IDRs flanking the central folded domain. (C) Length, motif content, and charge of compact RG regions. Each point represents one RG region, plotted as region length versus number of RG motifs. Color encodes mean net charge per region on a continuous scale. (D) RBP enrichment within the RG-region landscape. Left: net charge versus length for each RG region, with annotated RBPs highlighted. Right: number of RG motifs versus region length, with RBPs highlighted as in the left panel.

To translate this compositional enrichment into an operational, spacing-based rule, we quantified the organization of RG motifs using a kernel density estimation (KDE)-based contour approach adapted from our previous analyses of RG/RGG sequence grammar [49]. Curated RG/RGG proteins occupy a compact motif-spacing regime relative to generic disordered regions, motivating a short-linker criterion for defining RG tracts (Fig. 3B). Complementary summaries of inter-RG spacing are presented as cumulative distributions in Supplementary Fig. S3B, with corresponding linker-length values provided in Supplementary Table S4 (sheets S4C and S4E). Guided by these spacing statistics, we defined RG tracts as sequences in which consecutive RG motifs are separated by ≤7 residues, capturing the characteristic local density observed in curated RG/RGG proteins [49]. To avoid artificial fragmentation of otherwise cohesive clusters, we permitted linkers up to 15 residues only when bridging two compact tracts separated by a single longer spacer. Each resulting combined tract was then extended by five residues at both termini to preserve the surrounding sequence context (Fig. 3B, top) resulting in a comprehensible arginine-glycine-rich region. A representative example such a region from THOC4/ALYREF illustrates this segmentation strategy, with compact RG regions identified in both terminal IDRs flanking the central folded RNA-binding domain (Fig. 3B, bottom).

Applying this rule-based definition across the human proteome identified 1,008 compact RG regions across 823 proteins. Of these, 380 regions were found in 289 annotated RBPs. Within this catalogue, RG region length scaled with RG motif count, and most regions exhibited net positive charge (Fig. 3C; charge density summaries in Supplementary Fig. S3C, most regions are positive i.e. have positive net charge per residue value; per-region values in Supplementary Table S4, sheet S4A). While RBP-derived RG regions largely overlapped the global distribution, they were enriched for RG-dense and positively charged segments (Fig. 3D; relationship between charge density and RG density in Supplementary Fig. S3D, higher RG density might indicate higher charge in the region; RBP region identifiers in Supplementary Table S4, sheet S4F). Together, these analyses provide a proteome-wide catalogue of compact, charge-dense RG regions that complements the R-TAD dataset and enables systematic pairing of acidic activation domains with arginine-glycine-rich segments in subsequent interaction analyses.

### Electrostatic complementarity and charge patterning shape the simulated interaction landscape of R-TAD–RGG pairs

To map the sequence determinants of interactions between transcription-factor R-TADs and RBP RG/RGG regions, we carried out residue-level coarse-grained simulations using CALVADOS. In these simulations, each amino acid is represented by a single bead in implicit solvent, and interactions are governed by electrostatics together with residue-type–specific short-range ‘stickiness’ parameters tuned for intrinsically disordered proteins (Methods). From the resulting pairwise radial distribution functions we estimated second virial coefficients (B_22_) as a quantitative measure of effective attraction between segments. Due to the computational intractability of exhaustively sampling all 230 predicted transcriptional activation domains (R-TADs) and 1,008 RG/RGG regions (>2.3 × 10□ potential combinations), we assembled a representative simulation set using a staged, feature-guided strategy. A pilot matrix of five R-TADs and five RG/RGG segments spanning extremes and midpoints of length and net charge was first simulated in all 25 combinations to identify informative regions of feature space. Pilot correlations indicated comparatively stronger attraction for pairs with non-negative charge complementarity (a net-charge–based measure that is positive when the two segments carry opposite net charges; Methods), non-neutral mean pair charge (average NCPR outside –0.05 to 0.05), and average λ outside an intermediate band (0.45–0.56; Fig. S4A–C). These criteria were used to enrich subsequent sampling while retaining counter-examples outside these bounds, and candidates were additionally subsampled to maximize length diversity and limit redundancy. In a later expanded simulation stage, the opposite-charge number (OCN; a count-based measure of how many oppositely charged residues are available across the two segments; Methods) emerged as the strongest correlate of interaction strength, motivating subsequent rounds designed to improve electrostatic coverage via OCN-binned sampling (Fig. S4D). The resulting panel of 917 R-TAD–RGG pairs was simulated using the coarse-grained CALVADOS v2 model [42] at physiological ionic strength (150 mM NaCl, pH 7.0), and the complete list of simulated pairs, sequence features, and simulation outputs is provided in Table S5.

For each R-TAD–RGG pair, we computed pairwise radial distribution functions, g(r), and estimated second virial coefficients (B_22_) by integrating g(r) after applying a finite-size correction (Methods). B_22_ provides a dilute-solution measure of net intermolecular interactions derived from the radial distribution function. In this convention, more negative B_22_ indicates stronger effective attraction between segments (i.e., an increased probability of finding the two chains in close proximity relative to random mixing), whereas values near zero indicate near-ideal mixing. For convenience, when relating interaction strength to sequence features, we report log_10_(−B_22_) for attractive pairs (B_22_ < 0). Across the simulation panel, B_22_ values spanned nearly two orders of magnitude, ranging from near-ideal mixing to strongly attractive interactions. The overall distribution was unimodal, with a pronounced tail toward more negative B_22_ values, indicating a broad continuum of interaction strengths (Fig. 4A; Table S5). Representative g(r) profiles highlight this diversity. For example, a strongly interacting pair (SYNCRIP RGG/ FOXO6 R-TAD) exhibits pronounced short-range structure with a peak at 2-3 nm, consistent with frequent association (Fig. 4B, top). In contrast, a weakly interacting pair (FZD9 RGG/ ZNF165 R-TAD) shows g(r) close to unity, indicating minimal deviation from random mixing (Fig. 4B, bottom).

**Figure 4.**
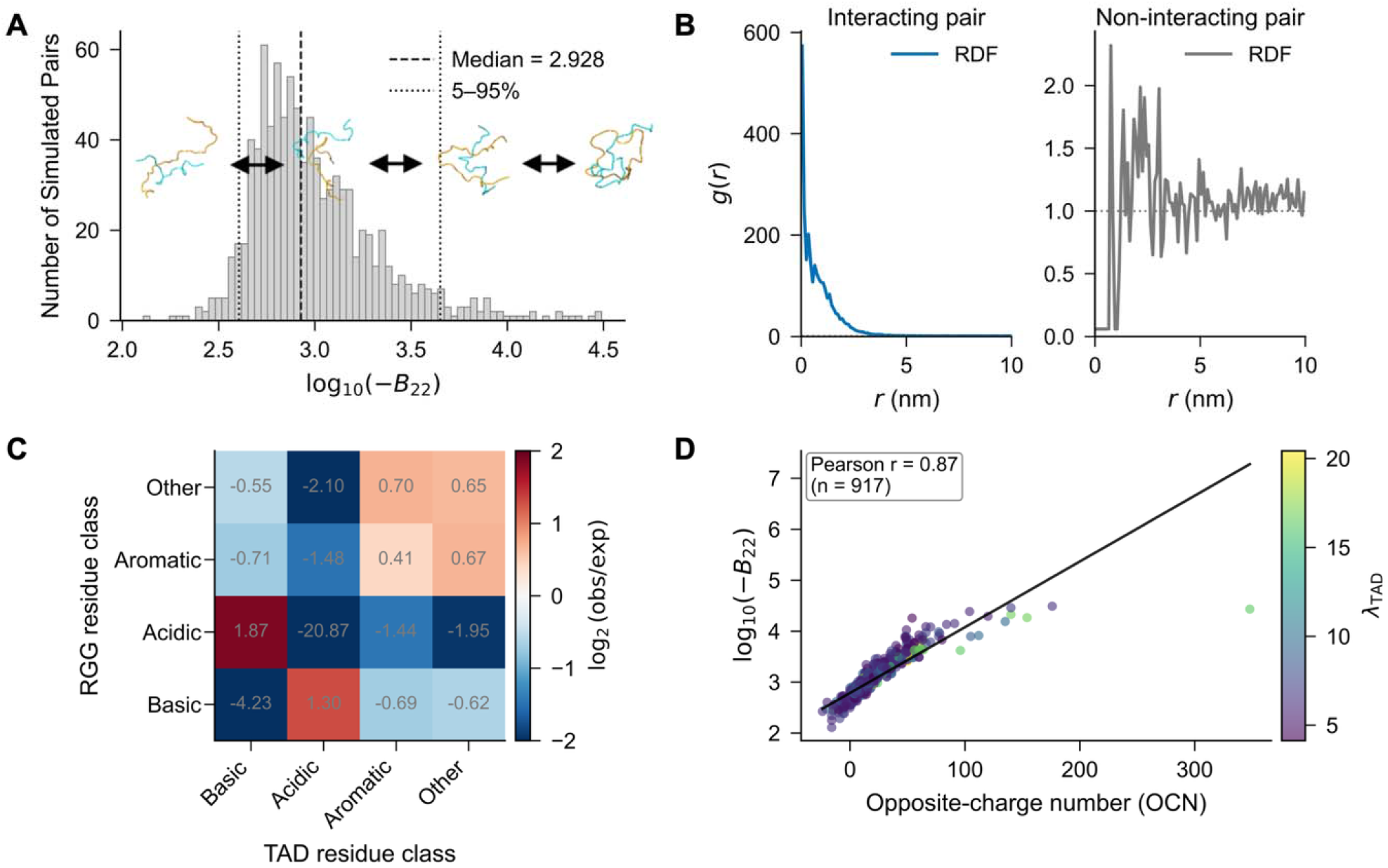
Simulated R-TAD–RGG interaction landscape highlights electrostatic complementarity as a dominant determinant of association strength. (A) Distribution of second virial coefficients (B_22_) estimated from CALVADOS v2 simulations of 917 R-TAD–RGG pairs at physiological ionic strength (150 mM NaCl). A finite-size correction was applied to g(r) prior to B_22_ integration (Methods). Vertical lines indicate the median and 5th–95th percentile range. Cartoon snapshots illustrate representative conformations across the weak-to-strong interaction continuum. (B) Representative radial distribution functions g(r) for an interacting and a non-interacting pair. The SYNCRIP RGG–FOXO6 R-TAD pair shows elevated short-range g(r), consistent with frequent association, whereas the FZD9 RGG– ZNF165 R-TAD pair remains near g(r) ≈ 1, indicating near-ideal mixing. (C) Residue-class contact enrichment across all simulated pairs, reported as log_2_(observed/expected) for contacts between residue classes in RGG regions (rows) and R-TADs (columns). Expected frequencies are derived from the residue-class composition of each partner. Enrichment highlights preferential contacts between basic (RGG) and acidic (R-TAD) classes relative to random expectation. (D) Opposite-charge number is strongly associated with simulated interaction strength. Scatter plot of log_10_(–B_22_) versus opposite-charge number (OCN) for 917 attractive pairs (B_22_ < 0). Points are colored by λ_R-TAD_; the black line denotes a linear fit, and the inset reports Pearson’s r.

To investigate the molecular basis of these interactions while controlling for compositional biases, we computed a residue-class contact enrichment map across all simulations, reported as log_2_(observed/expected) relative to the residue-class composition of each partner (Fig. 4C). This analysis revealed strong enrichment of contacts between basic residues in RG-rich regions and acidic residues in R-TADs, whereas other class combinations were depleted or near-neutral, supporting electrostatic complementarity as a dominant contributor to association under the simulated conditions. Consistent with this interpretation, the opposite-charge number (OCN) showed a strong linear correlation with simulated interaction strength when expressed as log_10_(–B_22_) (Fig. 4D; Pearson r = 0.87, n = 917). Across 917 simulated pairs, interaction strength increased with opposite-charge number over a broad dynamic range, consistent with electrostatic complementarity as a dominant contributor under the simulated conditions. Point color encodes λ_*R-TAD*_, although any additional contribution of R-TAD hydropathy is less visually apparent than the dominant dependence on opposite-charge number. Because the simulation panel was assembled using feature-guided sampling enriched for electrostatic complementarity, simple correlations between OCN and interaction strength may still reflect co-varying sequence properties, including length and hydropathy. Summary correlations between B_22_ and the full set of sequence and interaction descriptors used throughout this study are reported in Table S5.

### Sequence features dominated by opposite-charge number accurately predict TAD–RGG interaction strength

To extend insights from the simulated R-TAD–RGG interaction landscape to the full combinatorial sequence space, we developed a hybrid machine learning framework to predict interaction strength, defined as *y* = *log*_10_ (-*B*_22_) directly from sequence-derived descriptors (Fig. 5A; Table S6). For each R-TAD and RGG segment, we computed 22 features reflecting electrostatics, hydropathy, and CALVADOS λ-stickiness. This stickiness is a residue-specific parameter that sets the depth/strength of the short-range non-electrostatic interaction term, with pairwise strengths obtained by combining the λ values of the two interacting residues [49] (see Methods for details). Using these descriptors, we trained an ensemble of five independently initialized multilayer perceptron (MLP) regressors. The ensemble mean provided a robust baseline prediction *ŷ*_ens_. However, simulations of highly charge-complementary pairs entered a plateau regime that was not captured by *ŷ* (Fig. 5A; Fig. S5A). To address this, we trained a secondary residual model on the high-OCN subset (OCN ≥ 40) to predict the residual error *Δ*= *y*_sim_ - *ŷ*_ens_. This correction was down-weighted (residual weight ≤ 0.6), softly gated as a function of OCN to ensure a smooth transition, and capped at |Δ| ≤ 0.128 to prevent over-correction (Fig. 5A; Fig. S5B). The final hybrid model added this controlled residual term to the ensemble output, yielding a targeted adjustment that is negligible at low OCN but increases in the high-OCN regime (Fig. S5B), and re-centers prediction residuals toward zero when stratifying pairs into low- and high-OCN regimes (Fig. S5D).

**Figure 5.**
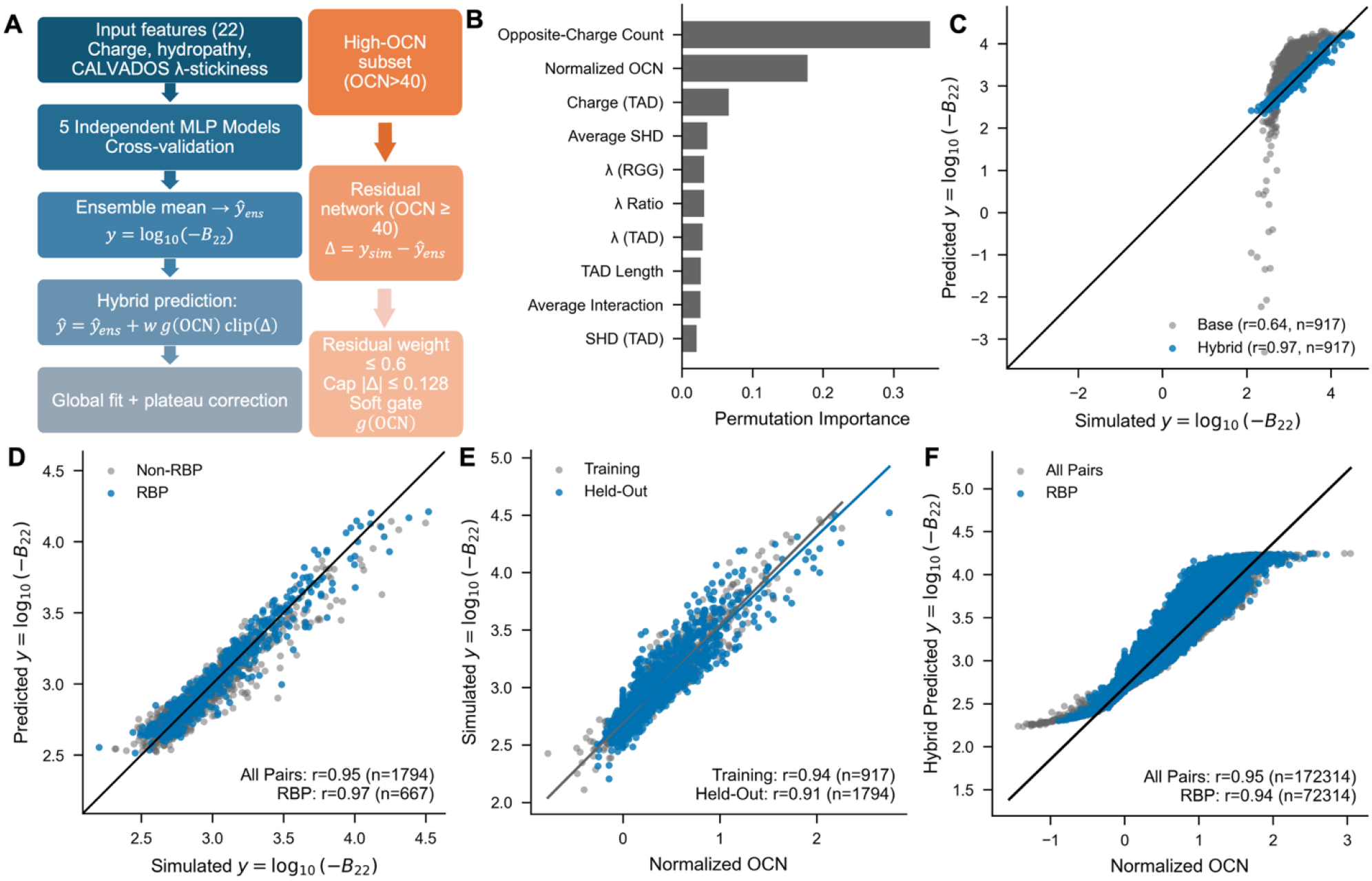
Hybrid machine learning model predicts TAD–RGG interaction strengths from sequence features. (A) Schematic of the hybrid modeling pipeline. For each R-TAD–RGG pair, 22 descriptors capturing electrostatics, hydropathy, and CALVADOS λ-stickiness are computed and used to train an ensemble of five MLP regressors. The ensemble mean provides a global prediction *ŷ*ens for interaction strength, defined as *y* = *log*_10_ (− *B*_22_) For highly charge-complementary pairs (OCN ≥ 40), a residual network is trained to model the remaining error *Δ* = y_sim_ − *ŷ*. The residual is softly gated by ONC, down-weighted (residual weight ≤ 0.6), and capped at | *Δ*| ≤ 0.128 before being added to *ŷ*_ens_, ensuring a smooth transition into the plateau regime. These plateau-aware parameters were selected empirically to improve fit in the high-OCN regime. (B) Permutation importance of the top features (held-out set). Opposite-charge count and normalized OCN dominate model importance, with additional contributions from R-TAD net charge, hydropathy patterning (average SHD), λ-stickiness terms (R-TAD, RGG, and their ratio), R-TAD length, and a coarse-grained interaction proxy. (C) Training-set performance comparing a baseline MLP (gray) and the hybrid model (blue). The baseline compresses strong interactions (Pearson *r* = 0.64, *n* = 917), whereas the hybrid closely matches simulations across the full range (Pearson *r* = 0.97). (D) Performance of the hybrid model on an independent sequence-exclusive held-out set (*n* = 1,794). Predicted versus simulated *y* = *log*_10_ (-*B*_22_) values follow the identity line (Pearson *r* = 0.95); RBP-containing pairs (blue) align with the global trend. (E) Simulated scaling of interaction strength with normalized OCN for training (gray) and held-out (blue) sets. Interaction strength increases monotonically and saturates at high charge complementarity (training *r* = 0.94, *n* = 917; held-out *r* = 0.91, *n* = 1,794). (F) Hybrid-predicted interaction landscape across all 207 × 930 combinations as a function of normalized OCN (gray; *r* = 0.95), with RBP-containing combinations highlighted (blue; *r* = 0.94). Interaction strength increases with normalized OCN and plateaus at high charge complementarity.

Permutation-importance analysis on the held-out set revealed that electrostatic features dominate prediction accuracy (Fig. 5B). The most informative variables were opposite-charge count and normalized OCN, followed by R-TAD net charge and secondary terms capturing hydropathy and stickiness (e.g., average SHD, *λ* of RGG and R-TAD, and their ratio), as well as R-TAD length and a coarse-grained mean interaction term (Fig. 5B; Table S6). Consistent with its plateau-aware design, the hybrid model expanded the dynamic range of predictions relative to a baseline MLP trained on the same features (Fig. 5C; Fig. S5C), consistent with reduced systematic underprediction in the high-OCN regime as summarized by residuals (Fig. S5D). The residual-gating threshold, residual weight, and cap were selected empirically to improve performance in the high-OCN regime; ablation and out-of-distribution tests will be required to establish robustness across sequence regimes and solution conditions. On the training set, the baseline MLP compressed strong interactions (Pearson *r* = 0.64, *n* = 917), whereas the hybrid closely ely followed the identity line (Pearson *r* =0.97; Fig. 5C). On an independent held-ou (*n* =1,794), hybrid predictions remained highly accurate (Pearson *r*= 0.97), and RBP-containing pairs followed the same predicted–simulated relationship as other combinations (Fig. 5D). There was no overlap of sequences betweeen the training and the held-out sets of TADs and RGGs.

Scaling analyses confirmed that interaction strength increases monotonically with normalized OCN before plateauing at high charge complementarity. This trend was observed in both the training and held-out simulations (Pearson *r* = 0.94 and *r* = 0.91, respectively; Fig. 5E), and was preserved when extending hybrid predictions to the full 207 × 930 combination space (Pearson *r* = 0.95), with RBP-containing combinations tracking the global pattern (Pearson *r* = 0.94; Fig. 5F). These findings indicate that the charge-driven grammar governing simulated interactions extends smoothly across the full R-TAD–RGG sequence space within the present simulation framework, with RBPs spanning the full interaction range rather than forming a distinct outlier class. In addition, independent FINCHES-derived interaction scores [50] correlate with simulated interaction strength in the training set and with hybrid-predicted interaction strength across all pairs (Fig. S6A–B; Table S7), supporting the same overall trend within an independent sequence-based scoring framework.

### NMR titrations reveal a hierarchy of TAD–RGG interaction strengths consistent with sequence-based predictions

To evaluate whether sequence-based interaction predictions reflect relative binding strengths, we performed NMR HSQC titration experiments using five acidic R-TADs: ATF4, FOXO4, FOXO6, TFE3, and LEF1, against four RG/RGG-rich partners: the N- and C-terminal fragments of THOC4, G3BP1 RGG, and TAF15. These interaction pairs were selected to span a range of predicted affinities and to sample distinct titration behaviors. Each titration was classified into one of three regimes based on the spectral response: (i) K_d_ fit, characterized by progressive chemical shift perturbations (CSPs) that could be quantitatively fitted using TITAN under the applied model; (ii) line broadening, where peak intensity loss and disappearance prevented robust extraction of dissociation constants under the tested conditions; and (iii) no binding (NB), defined by negligible spectral changes even at high ligand excess. The classification outcomes and fitted parameters are summarized in Figure 6A and Table S8, and representative ^1^H,^1^□N HSQC spectra illustrating K_d_-fit and line-broadening behaviors are shown in Figure 6B. LB titrations were therefore interpreted qualitatively and were not assigned K_d_ values. The classification outcomes and fitted parameters are summarized in Figure 6A and Table S8. Quantitative TITAN analyses yielded apparent dissociation constants in the high micromolar range for multiple interactions involving THOC4 and G3BP1. The FOXO4–THOC4 N-terminal interaction exhibited the strongest affinity (K_d_ = 48.5 ± 1.6 µM), followed by FOXO6–THOC4 N-term (K_d_ = 67.2 ± 2.2 µM), ATF4–THOC4 N-term (K_d_ = 183.7 ± 6.6 µM), and TFE3–THOC4 N-term (K_d_ = 250.8 ± 41.9 µM). In contrast, titrations involving TAF15 primarily exhibited line broadening, consistent with a different exchange regime under the tested conditions and precluding quantitative K_d_ extraction, while LEF1 failed to show detectable binding with any partner under these conditions.

**Figure 6.**
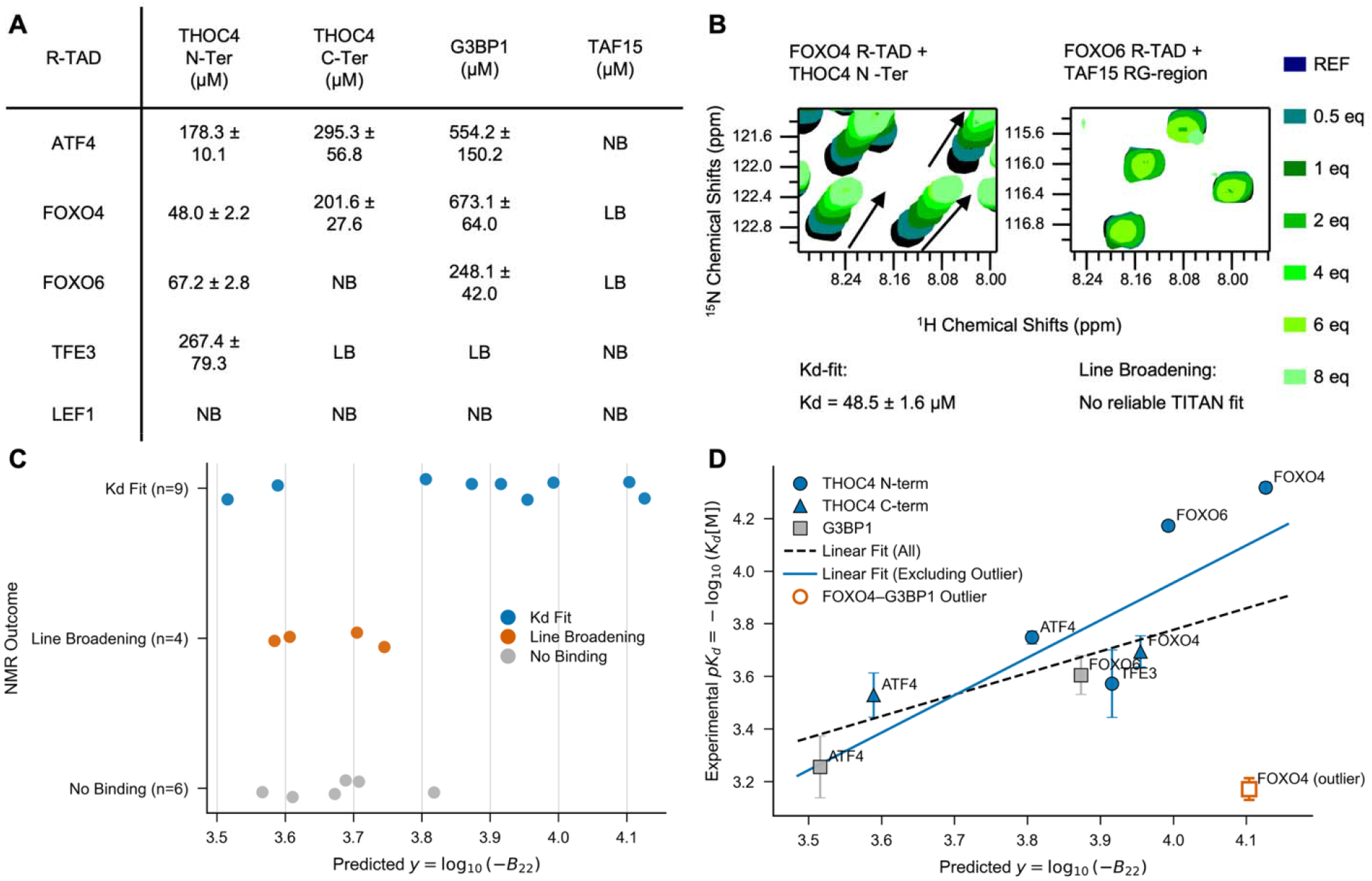
NMR titrations validate predicted R-TAD–RGG interaction strengths. (A) Summary table of [^15^N]–HSQC titrations between five R-TADs (ATF4, FOXO4, FOXO6, TFE3 and LEF1) and four RG/RGG partners (THOC4 N-term, THOC4 C-term, G3BP1 RGG and TAF15). Where progressive CSPs enabled quantitative analysis, dissociation constants are reported as K_d_ (µM ± TITAN fit error). Titrations that could not be robustly fit are annotated as LB (line broadening) or NB (no binding). (B) Predicted interaction strength y = *log*_10_ (-*B*_22_) for each tested pair stratified by experimental outcome (K_d_ fit, line broadening, no binding). (C) Relationship between predicted interaction strength and experimental affinity for the subset of interactions with TITAN-derived K_d_ values, plotted as *pK*_*d*_ = - *log*_10_ (*K*_*d*_[*M*])) with propagated errors. FOXO4–G3BP1 deviates from the overall trend (annotated as an outlier); linear fits are shown for all K_d_-fit points and for the dataset excluding FOXO4–G3BP1. (D) Representative HSQC zoom regions illustrating distinct titration behaviours. Left: a K_d_-fit example showing smooth, trackable CSPs across increasing equivalents. Right: a line-broadening example where intensity loss/peak disappearance prevents reliable fitting under the current conditions. Colours indicate increasing equivalents.

Experimental outcomes stratified consistently with predicted interaction strength, expressed as *y* = *log*_10_ (-*B*_22_): interactions yielding quantitative K_d_ fits were enriched at higher predicted values, non-binding pairs clustered at lower values, and line-broadening cases spanned intermediate-to-high predicted interactions strengths (Figure 6C). For LB cases without extractable K_d_ values, qualitative titration behavior aligned with the predicted interaction hierarchy. Among interactions yielding quantitative fits (n = 9), apparent experimental affinity (expressed as *pK*_*d*_ = - *log*_10_ (*K*_*d*_[*M*])) correlated positively with predicted interaction strength (Figure 6D). One outlier, FOXO4–G3BP1, deviated markedly from this trend: excluding this point improved concordance, indicating that the predictive power of the sequence-derived model for the remaining interactions. Moreover, simulation-derived interaction strengths for the same set recapitulated both the quantitative affinity trends and categorical stratification of titration outcomes (Figure S7; Table S8), reinforcing that sequence-based predictions capture key features of the underlying biophysical interaction landscape under the tested conditions.

## Discussion

The integrative analyses presented here reveal that interactions between transcription factors and RNA-binding proteins are not isolated curiosities but pervasive features of gene regulation. Network-level mapping shows that RBPs are disproportionately connected to TFs relative to other protein classes in curated interaction datasets, highlighting a broad interface within transcriptional regulatory circuits and linking DNA- and RNA-based processes. Notably, this connectivity is heterogeneous across the TF repertoire: while many TFs exhibit modest RBP connectivity, a substantial subset is strongly RBP-enriched, with RBPs comprising more than half of their interaction partners (Fig. 1B). Network centrality analysis further suggests that this connectivity is concentrated in a cohort of high-betweenness of TF and RBP hubs, positioning these proteins to bridge transcriptional regulation with RNA-processing and signaling subnetworks (Fig. 1C–D). These observations place the TF–RBP interface along a functional continuum of gene-expression pathways, spanning transcription initiation, mRNA processing, translation, and stress-induced condensate dynamics. RGG-containing RBPs, in particular, have been implicated in chromatin remodeling, pre-mRNA splicing, mRNA export, translation, and even DNA repair [51], while hnRNPA2/B1 functions in transcription, splicing, RNA transport, and translational regulation [52]. This multifunctionality is exemplified by the FET family proteins (FUS, EWS, and TAF15), which bind both RNA and DNA, participate in transcription, splicing, and microRNA processing, and relocalize to stress granules, linking transcriptional and post-transcriptional regulation to stress-responsive translation control [51].

Within this regulatory landscape, acidic/hydrophobic transactivation domains and positively charged RG/RGG motifs appear to form complementary disordered “grammars.” Building on this observation, we defined RG/RGG-binding activation domains based on linker length rules between consecutive RG repeats from [52], which captures the high local RG density and short inter-motif spacing characteristic of RBP low-complexity regions. Binding to this peptide identified a small set of robust interactors (“seed R-TADs”), which we extended by incorporating homologous acidic segments from related TF families. Comparative feature analysis showed that these R-TADs are not generic acidic regions, but short disordered segments enriched in aromatic residues and defined by dispersed charge and low-complexity sequence patterns. However, these associations do not by themselves establish causal necessity or sufficiency for RG/RGG binding. In particular, mutational perturbation of TAD sequence features and composition- or dipeptide-preserving null models will be needed to disentangle the relative contributions of composition and patterning. Leveraging this grammar, we scaled predictions to the full TF repertoire using an ensemble classifier trained on sliding windows across TF sequences, identifying 230 predicted R-TADs across 190 TFs. These predictions overlap incompletely with previously published AD/TAD catalogues, supporting the view that an RG/RGG-binding–oriented definition recovers established activation domains while nominating previously uncharacterized candidates for TF–RBP coupling. In parallel, we established a spacing-based definition of RG regions by quantifying motif organization and applying an explicit linker rule to identify compact RG-rich regions. Applied proteome-wide, this yielded 1,008 compact RG regions across 823 proteins, including 380 in annotated RBPs. These sequences are characterized by high RG motif density and net positive charge, defining a standardized “RG region” that complements the R-TAD catalogue and supports systematic pairing analyses. For example, ALYREF (THOC4) contains such regions and has been implicated as a multifunctional regulator at the interface of transcription and RNA export.

By quantifying R-TAD–RGG association using second virial coefficients (*B*_22_) derived from coarse-grained simulations, we move beyond qualitative interaction descriptions to define a comparative scale of interaction strength under controlled solution conditions. Across the simulated panel, interaction strength is strongly shaped by electrostatic complementarity: residue-class enrichment analyses show preferential contacts between basic residues in RG-rich regions and acidic residues in R-TADs, with the number of oppositely charged residues emerging as a dominant axis of interaction strength under the simulated conditions. Importantly, we moved beyond post hoc interpretation by developing a hybrid machine-learning model that predicts interaction strength, expressed as *y* = *log* _10_ (-*B*_22_), directly from sequence-derived features. Permutation-importance analysis demonstrated that electrostatic descriptors, particularly opposite-charge metrics, dominate predictive performance, while secondary contributions arise from hydropathy- and stickiness-related properties, including CALVADOS *λ*-based features, supporting a charge-dominated landscape modulated by additional physicochemical factors. Because the feature set, simulation framework, and sampling strategy were all designed to probe electrostatic complementarity, this agreement across analyses should be interpreted as support for an electrostatically dominated model under the present assumptions rather than as a complete exclusion of non-electrostatic contributions.

A notable outcome of this model is that it makes the structure of the interaction landscape explicit. The hybrid framework was designed to address a plateau in simulated affinity at high charge complementarity, which was not captured by a single global regressor. By incorporating a controlled residual model for high-OCN pairs, the model maintains accuracy across the full interaction range. Its strong performance on independent held-out data supports the conclusion that the sequence grammar learned from simulations extends beyond the simulated subset within the present modeling framework. When extended to the full combinatorial R-TAD–RGG space, predicted interaction strengths follow the same monotonic trend with normalized charge complementarity, plateauing at high OCN values. RBP-containing pairs track this global trend rather than forming a distinct subclass, suggesting that TF–RBP pairing is a general feature of IDR-based interaction space. In practical terms, the model enables principled prioritization of TF–RBP pairs for experimental validation and supports rational engineering or discovery of disordered modules with tailored association propensities.

NMR titrations provided an orthogonal experimental validation of the predicted interaction hierarchy. Across two acidic TADs and four RG/RGG partners selected to span the predicted affinity spectrum, the magnitude of HSQC perturbations and line broadening scaled with predicted strength: strongly predicted partners induced widespread chemical shift perturbations across the disordered TADs, while weakly predicted partners produced only minimal or localized effects (Fig. 6). These results support the interpretation that the model captures relative interaction strengths and that these contacts form a continuum of dynamic, multivalent complexes rather than discrete, structured interfaces. At the same time, because the validated pairs were selected to span the predicted interaction landscape, these experiments support the ranking capacity of the framework but do not independently exclude additional non-electrostatic determinants of binding.

Importantly, the goal of this study was not merely to catalogue TF–RBP interactions, but to infer generalizable principles of IDR crosstalk that can inform functional hypotheses. Translating interaction propensities into cellular outcomes, however, requires consideration of broader contextual factors, including macromolecular crowding, RNA binding, and regulatory post-translational modifications. In particular, arginine methylation, a known regulator of RG/RGG domains, is expected to modulate interactions without altering net charge, by changing steric properties, charge distribution, or hydrogen bonding potential [30]. Within these constraints, the convergence of network topology, sequence grammar, simulation-based energetics, predictive modeling, and experimental validation supports a model in which acidic activation domains and compact RG regions can function as modular, sequence-encoded interfaces linking transcription with RNA metabolism. More broadly, this work demonstrates that even fuzzy interactions between disordered regions can be mechanistically rationalized, and in many cases predicted, using a tractable set of physicochemical rules derived directly from sequence.

### Limitations of the study

This study combines curated interactome analysis, sequence-based identification of RG/RGG-binding transactivation domains (R-TADs), coarse-grained CALVADOS simulations, machine-learning prediction, and targeted NMR validation to define a quantitative framework for TF–RBP coupling through disordered regions. Several limitations should be considered. First, the simulation-derived interaction landscape was generated using a coarse-grained implicit-solvent model under fixed solution conditions and therefore does not capture the full molecular detail, sequence-context dependence, or cellular heterogeneity of TF–RBP interactions. Second, the simulated pair set was selected to sample informative regions of sequence space rather than exhaustively cover the full combinatorial landscape, so model behavior should be interpreted within the range represented by the present training data. Third, the predictive framework is trained against simulation-derived interaction strengths rather than direct biochemical affinity measurements for all pairs; accordingly, it is best viewed as a prioritization and ranking tool rather than a substitute for experimental determination of binding constants. Fourth, the NMR validation set is intentionally limited and supports the predicted interaction hierarchy, but it does not establish a complete biophysical description for all predicted partners or exclude additional contributions from RNA, post-translational modifications, multivalency, or cellular cofactors. Finally, network-level enrichment analyses are based on curated PPI resources and annotation-defined RBP sets, and therefore capture robust trends in available datasets rather than a degree-controlled estimate of class-specific binding preference.

## Materials and Methods

### Dataset collection

Protein sequences of the human proteome were downloaded from UniProt (release 2023_03). Lists of transcription factors (TFs) and RNA-binding proteins (RBPs) were obtained from curated datasets [43,44]. The RBP annotation used for network analyses was based on EuRBPDB, an annotation-based resource that includes both canonical RBPs containing known RNA-binding domains and non-canonical RBPs supported by RBPome datasets and literature curation. Because curated RBP resources can include multifunctional regulators, the RBP class in Figure 1 should be interpreted as an annotation-based reference set. Only canonical isoforms were used for downstream analysis. Intrinsically disordered regions (IDRs) were predicted using SPOT-Disorder2 [56], requiring a minimum disordered stretch of 30 residues.

### Identification of TADs and RG/RGG regions

Transactivation domains or TADs were predicted using a multi-classifier machine-learning framework [57,58] trained on experimentally verified acidic and hydrophobic activation domains from p53, FOXO, LEF1, and related TFs. Input comprised 15-residue fragments (sliding window) from disordered TF regions. For seed-level sequence profiling (Fig. 2B), experimentally defined and homologous R-TAD segments were compared to length-matched disordered windows from other TFs as a qualitative baseline. Composition- and dipeptide-preserving null models were not applied in this analysis. For proteome-wide prediction, each fragment was encoded by 143 molecular sequence features [48], including amino-acid composition, SLiMs, charge parameters (NCPR, κ*, SCD, SHD), aromaticity, and stickiness (λ) [48]. The scripts to derive these features are available at https://github.com/IPritisanac/idr.mol.feats. Positive labels were assigned to windows overlapping experimentally validated seed R-TADs and homologous acidic segments, whereas negative labels were assigned to length-matched windows from other disordered TF regions that did not overlap curated R-TADs. Classifiers (logistic regression and MLPs) were implemented in scikit-learn (v1.3) [56] with SMOTEENN resampling [57,58] and grid-search hyperparameter optimization. Models were evaluated on validation folds using F1, F0.5, and recall, and predictions from the best-performing models were averaged. Fragments predicted as TADs by all 140 models (consensus) were accepted as high-confidence TADs, and overlapping windows were merged into contiguous segments. This procedure yielded 230 predicted R-TADs across 190 TFs.

RG/RGG regions were identified using an expanded rule-based algorithm derived from the statistical distribution of RG motifs across disordered regions in the human proteome. Predicted IDRs from SPOT-Disorder2 were scanned for RG and RGG motifs using sliding windows of 20–100 amino acids. Motifs occurring within ≤ 7 residues were grouped into tracts, and adjacent tracts separated by ≤ 15 residues were merged into continuous regions. These empirically derived thresholds were obtained from proteome-wide analyses of inter-motif spacing and motif density. Each tract was extended by ±5 residues to preserve flanking sequence context and annotated with motif count, tract length, and disorder probability for downstream simulation and feature extraction.

### Simulation of TAD–RGG interactions

#### CALVADOS model

We performed residue-level coarse-grained molecular dynamics simulations using the CALVADOS v2 framework for intrinsically disordered proteins. In CALVADOS, each amino acid is represented by a single bead and the solvent is treated implicitly. Nonbonded interactions include electrostatics together with residue-type–specific short-range interactions that capture sequence-dependent association propensities of disordered regions [42].

#### System setup and conditions

Each simulation contained one R-TAD segment and one RG/RGG segment (1:1 stoichiometry) in a cubic periodic simulation box (side length 20 nm). Termini were charged on both ends (charge_termini = ‘both’). Systems were initialized using a grid-based placement procedure to avoid steric overlaps. Simulations were carried out at 298 K, pH 7.0, and physiological ionic strength (150 mM NaCl equivalent).

#### Integrator, ensemble, and sampling

Dynamics were propagated using Langevin dynamics under constant-volume (NVT) conditions. Langevin dynamics is a stochastic thermostat that maintains the target temperature by adding a friction term and random thermal forces to the equations of motion, providing stable temperature control in implicit-solvent simulations. We used a timestep of *Δ*t = 10 fs and simulated each replica for 100,000,000 integration steps (total 1 μs). Coordinates were saved every 10,000 steps (100 ps) for analysis, and checkpointing was enabled to allow robust restarts.

#### Replication strategy

To improve sampling and quantify variability, we ran five replicas per R-TAD–RGG pair. Replicas were initialized with distinct random seeds (and independent initial velocity assignments where applicable) to promote statistical independence across trajectories and reduce sensitivity to the starting configuration.

#### Simulation Protocol

Representative 917 TAD–RGG pairs were simulated using CALVADOS v2 [42] at 298 K and 150 mM ionic strength in cubic periodic boxes (20 nm side length) for 1 μs per replica (five replicas per pair). Dynamics were propagated using Langevin dynamics under constant-volume (NVT) conditions with a timestep of 10 fs (see simulation scripts for full settings). Replicas were initialized with distinct random seeds (and independent initial velocities where applicable) to promote statistical independence across trajectories and reduce sensitivity to the initial configuration (42)

### Radial distribution functions and B_22_ computation

Center-of-mass radial distribution functions g(r) between chains A and B were calculated using MDTraj and MDAnalysis (0.1 nm binning). Second virial coefficients (B_22_) were computed by integrating the radial distribution function g(r) according to the Kirkwood–Buff relation:

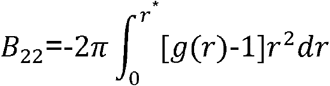

where *r*^*^= L/2 under periodic boundary conditions. Finite-size deviations of g(r) from unity near *r*^*^ were corrected following the procedure of [59], which builds on the Hummer tail correction [60]. The RDF tail was averaged over a window [*r*^*^ -Δ, *r*^*^] with *Δ* = 2 nm to obtain a background offset 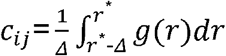. This averaged value replaced the ideal limit g(r→∞) = 1 in the integration, yielding the corrected form:

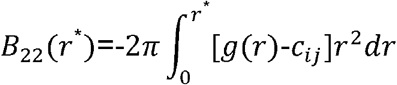

as implemented by [59] for sugar–protein RDFs. This approach minimizes artificial cubic divergence and improves convergence of B_22_ in finite simulation boxes.

### Sequence feature computation

Sequence-level descriptors were computed to capture electrostatics, hydropathy, and sequence patterning:

- CALVADOS parameters (λ, σ). In CALVADOS v2, each residue type is assigned a stickiness parameter λ that scales the strength of short-range, non-electrostatic interactions and an effective bead size σ that sets the interaction length scale in the coarse-grained potential (electrostatics treated separately). The λ and σ values used here are provided in residues.csv included with the code/data release.
Lambda Ratio

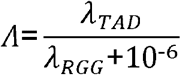
  - Net charge and charge-derived features Net charge of a segment was computed as 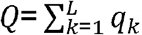 with residue charges *q*_*k*_ assigned according to our sequence model (D/E negative; K/R positive; histidine handled consistently with the pipeline). Net charge per residue was *NCPR*=*Q*/*L*, and absolute charge count was |*q*| = ∑_*k*_ |*q*_*k*_|.
- Charge complementarity was quantified as a net-charge summary metric

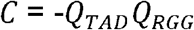

which is positive for oppositely charged pairs and negative for like-charged pairs. We additionally report the charge-balance metric

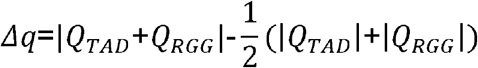

which measures cancellation of net charge upon pairing. Charge Asymmetry was defined as

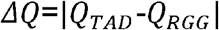
- Opposite Charge Number (OCN) We define OCN as the difference between the number of potentially oppositely charged residue pairs and the number of like-charged residue pairs available across the two segments, based on counts of acidic and basic residues (a composition-based electrostatic capacity metric that does not use residue positions).

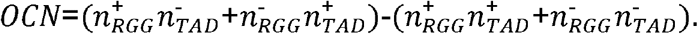

This definition increases with the availability of opposite charges while penalizing like-charge capacity, and thus captures net electrostatic pairing potential at the level of composition. Normalized OCN was defined as

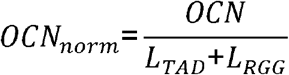
- Sequence charge decoration (SCD) [61] and sequence hydropathy decoration (SHD) [62] were computed as in the original definitions and applied separately to each segment. We report segment-level values (e.g., SCD_TAD, SHD_RGG) and the mean across the two segments (Average_SCD, Average_SHD) where indicated. (47)(46)
- Average λ and σ, aromatic fraction (Y/F/W), and the short-range hydropathy interaction integral *U*_*AH*_.
- Hydropathy Difference

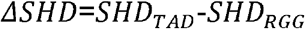

All features were computed in Python 3.11 using Numba for acceleration [63,64].

### Machine-learning framework

#### Model architecture

A multilayer perceptron (MLP) regression network was implemented in TensorFlow (v2.12) [65] and Keras [66] to predict

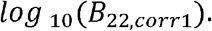

The network consisted of three fully connected hidden layers (100, 50, 25 neurons; tanh activation) and was trained with the Adam optimizer (learning rate = 0.01, *α*=0.01, 500 iterations).

Features were standardized using StandardScaler (scikit-learn v1.3) [56]. Training used early stopping and adaptive learning-rate reduction under 10-fold cross-validation.

#### Feature engineering

Four engineered descriptors were added to enhance scale-invariant and comparative biophysical interpretation:

1. Normalized OCN

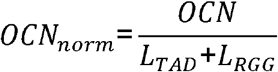
2. Charge Asymmetry

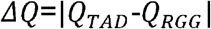
3. Hydropathy Difference

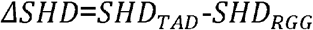
4. Lambda Ratio

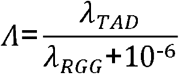

#### Ensemble training

To reduce variability from stochastic initialization and train/test splitting, five MLPs were trained independently using 75:25 train–test splits.

Final predictions were obtained by averaging the outputs of all five models.

#### Residual correction

A secondary dense neural network (DenseNN) was trained to correct systematic underestimation at high electrostatic imbalance.

Residuals were computed as

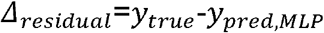

for samples with *OCN*>40.

The final hybrid predictor was:

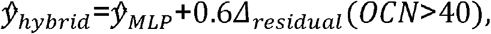

with the correction amplitude capped at ±0.128.

This module was embedded within the TensorFlow computation graph for batched inference.

#### Feature selection

Permutation importance (10 random shuffles per feature) quantified individual feature contributions via changes in

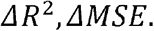

#### Evaluation and reproducibility

Model performance was quantified using MSE and R^2^ across cross-validation, test, and external evaluation sets.

All trained MLPs, scalers, and preprocessing configurations were serialized as .pkl bundles.

All computations were executed on HPC CPU nodes.

#### Plasmids

Expression constructs for all sequences* were generated by synthesis of the corresponding optimized cDNAs and insertion into pETM11 vector containing an N-terminal His_6_–Protein A tag using NcoI/BamHI restriction sites for Escherichia coli (E. coli) expression (Genscript).

* the sequences include

1. human FOXO4 461-505 (Uniprot ID P98177)
2. human LEF1 1-101 (Uniprot ID Q9UJU2)
3. human FOXM1 685-801 (Uniprot ID Q08050)
4. human p53 1-94 (Uniprot ID P04637)
5. human TFE3 475-575 (Uniprot ID P19532)
6. human ATF4 41-141 (Uniprot ID P18848)
7. human FOXO6 418-472 (Uniprot ID A8MYZ6)
8. human THOC4 (Uniprot ID Q86V81) full length and the shorter fragments-residues 1-100 (THOC4 N terminus) and residues 182-257 (THOC4 C terminus)
9. human TAF15 521-592 (Uniprot ID Q92804)
10. human G3BP1 421-466 (Uniprot ID Q13283)

#### Protein expression and purification

For expression of recombinant unlabeled or uniformly ^15^N-labeled proteins, the pETM11 His_6_–Protein A constructs were transformed into *Escherichia coli* BL21(DE3) Star cells. 10ml of liquid preculture were then transferred in 1L lysogeny broth (LB) medium or in case of isotope labeled in minimal medium supplemented with 6 g of 12C6H12O6 (Roth) and either 3 g of 14NH4Cl (Roth) or 1g of 15NH4Cl (Sigma). Cells were grown to an optical density (OD) (600nm) of 0.8 and protein expression was induced by addition of 0.5 mM IPTG at 20°C for 16 hours. Cells pellets corresponding for protein expression of unlabeled or 15N labeled disordered proteins fragments were harvested and sonicated in denaturating lysis buffer (50mM Tris-HCl pH 7.5, 150mM NaCl, 20mM Imidazole, 6M urea). ZZ-His6 recombinant proteins were then purified using Ni-NTA agarose (Qiagen) and the ZZ-His6 tag was cleaved with TEV protease treatment. After desalting to a low imidazole buffer (50mM Tris pH7.5, 150mM NaCl, 20mM Imidazole, 2mM TCEP). The untagged proteins were then isolated performing a second affinity purification using Ni-NTA beads. On the other hand, positively charged fragment-THOC4 1-100 were then subjected to a heparin column (Cytiva) to separate the cleaved protein from the tag. A final exclusion chromatography purification step was performed in the final buffer on a gel filtration column (Superdex 300 increase, Cytiva). Protein concentrations were estimated from their absorbance at 280 nm, assuming that the ε at 280 nm was equal to the theoretical ε value. Final buffer for all validation experiments was 20mM HEPES, 150mM NaCl, 2mM TCEP, pH 6.5, with proteins at 25μM or 50μM concentration for the reference experiments.

#### Experimental validation by NMR

^1^H,^1^□N HSQC titration experiments were performed at 298 K using uniformly ^1^□N-labeled proteins in 20 mM HEPES, 150 mM NaCl, 2 mM TCEP, pH 6.5. Increasing amounts of unlabeled binding partner were added to the labeled protein at molar ratios of 0.5, 1, 2, 4, 6, and 8 equivalents. Spectra were recorded on a 600 MHz Bruker Avance Neo spectrometer equipped with a TXI 600S3 probehead, processed using Topspin and NMRPipe [67]. CcpNmr Analysis 2.5 [68] was used for titration figures. Chemical shift perturbations (CSPs) were calculated as

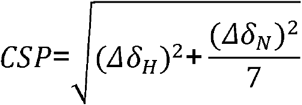

Resonances showing well-defined chemical shift perturbations and minimal overlap were selected for analysis. NMR lineshape fitting was performed in TITAN v1.6 [69]. Although the interaction likely involves a heterogeneous ensemble of bound conformations, as expected for intrinsically disordered proteins, these states are averaged on the NMR chemical shift timescale, allowing the titration data to be treated as an effective two-state process (free ⇌ bound) for global fitting. Dissociation constants (K_d_) were robustly determined from the global fits, whereas dissociation rate constants (koff) could not be reliably constrained because of fast exchange on the chemical shift timescale, as reflected by large bootstrap (100 replicas) uncertainties. Accordingly, only K_d_ values are reported and discussed.

#### Data analysis and visualization

All analyses were performed in Python 3.10 with NumPy, pandas, Matplotlib, and seaborn. Statistical evaluations employed Pearson correlation and 10,000-fold bootstrap resampling.

Code used for this study is publicly available on Zenodo and Github. The TAD prediction pipeline is available as idr-sequence-predict (DOI: 10.5281/zenodo.19080352), with its associated dataset deposited separately (DOI: 10.5281/zenodo.19080226). CALVADOS-based IDR–IDR interaction analysis workflows are available as idr-interaction-analysis (DOI: 10.5281/zenodo.19485640). The sequence-based interaction prediction workflow trained on simulation-derived B22 values is available as idr-interaction-predict (DOI: 10.5281/zenodo.19485655). Scripts and source-data workflows used to generate the figures in this manuscript are available as rtads-rgg-interaction (DOI: 10.5281/zenodo.19485560).

Processed supporting data associated with this study are deposited on Zenodo (DOI: 10.5281/zenodo.19485134), including analysis scripts, input files, processed datasets, radial distribution function files, per-pair energy pickle files, taken-out validation data, prediction-related resources, combined trajectories, and NMR spectra.

## Acknowledgements

We thank members of the Madl and Lindorff-Larsen laboratories for discussions and feedback. The authors acknowledge computational resources provided by the Danish e-Infrastructure Consortium (DeiC) through national HPC infrastructure at the University of Copenhagen, and by the MedBioNode cluster at the Medical University of Graz. We also acknowledge the use of Zenodo for code and data deposition. Y.K. and A.R. were trained within the framework of the PhD program Biomolecular Structures and Interactions (BioMolStruct). S.U. was trained within the framework of the PhD program Metabolic and Cardiovascular Disease (DK-MCD). For open access purposes, the author has applied a CC BY public copyright license to any author accepted manuscript version arising from this submission. T.M. thanks the Center for Medical Research, Medical University of Graz, Graz, Austria for laboratory access. T.M. is grateful to the Austrian Science Fund (FWF) for excellence cluster 10.55776/COE14, Grants DOI 10.55776/P28854, 10.55776/I3792, 10.55776/DOC130, and 10.55776/W1226, the Austrian Research Promotion Agency (FFG) grants 864690 and 870454; the Integrative Metabolism Research Center Graz; the Austrian Infrastructure Program 2016/2017; the Styrian Government (MetAGE, Zukunftsfonds, doc.fund program); the City of Graz (MetAGE, doc.fund); and BioTechMed-Graz (flagship project). This project was funded in part by the FFG and the European Union (EFRE) under grant 912192.

## Author contributions

Y.K. conceived and performed computational analyses, developed the sequence-analysis and prediction workflows, carried out simulations and downstream modeling, analyzed the data, did the preliminary experimental work and wrote the manuscript.

A.H. contributed to experimental work, analysis and writing.

J.P. contributed to experimental work and analysis.

S.U. performed experimental work.

A.R. contributed to analysis of experimental work.

I.P. contributed to conceptual input and analysis.

S.v.B. supervised and contributed to simulation strategy, analysis, and manuscript editing.

K.L.-L. supervised computational aspects of the work and contributed to interpretation and manuscript editing.

T.M. conceived and supervised the study, contributed to data interpretation, and edited the manuscript.

