## Supplementary figures and legends for "Proteome-wide identification and modeling of interactions between transactivation domains and arginine-glycine-rich regions"

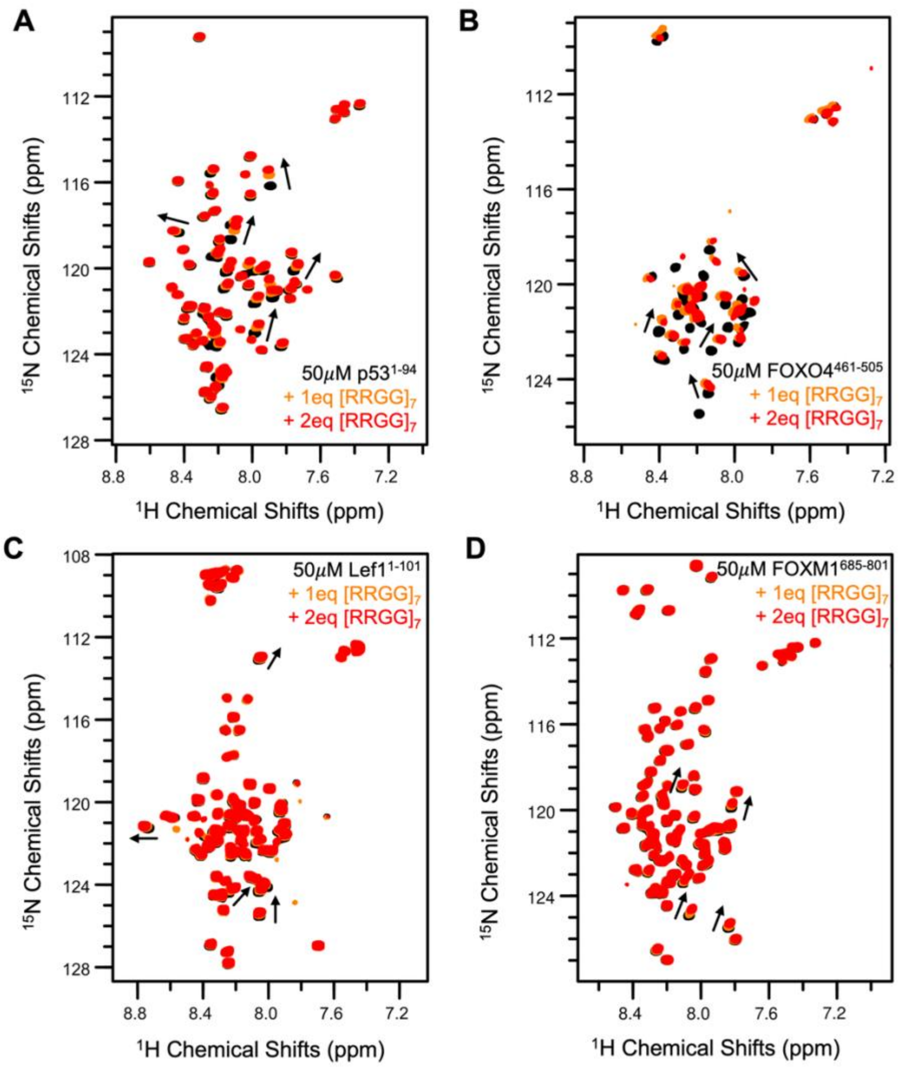


*Figure S1 | Selection of seed R-TADs based on positive interaction with [RRGG]$_7$ peptide*

(A) 50$\mu M$ p53 1-94 region (black) titrated with 1 (orange) and 2 (red) equivalents of [RRGG]$_7$ peptide, major shifts marked with black arrows.
(B) 50$\mu M$ FOXO4 461-505 region (black) titrated with 1 (orange) and 2 (red) equivalents of [RRGG]$_7$ peptide, major shifts marked with black arrows.
(C) 50$\mu M$ Lef1 1-101 region (black) titrated with 1 (orange) and 2 (red) equivalents of [RRGG]$_7$ peptide, major shifts marked with black arrows.
(D) 50$\mu M$ FOXM1 685-801 region (black) titrated with 1 (orange) and 2 (red) equivalents of [RRGG]$_7$ peptide, major shifts marked with black arrows.

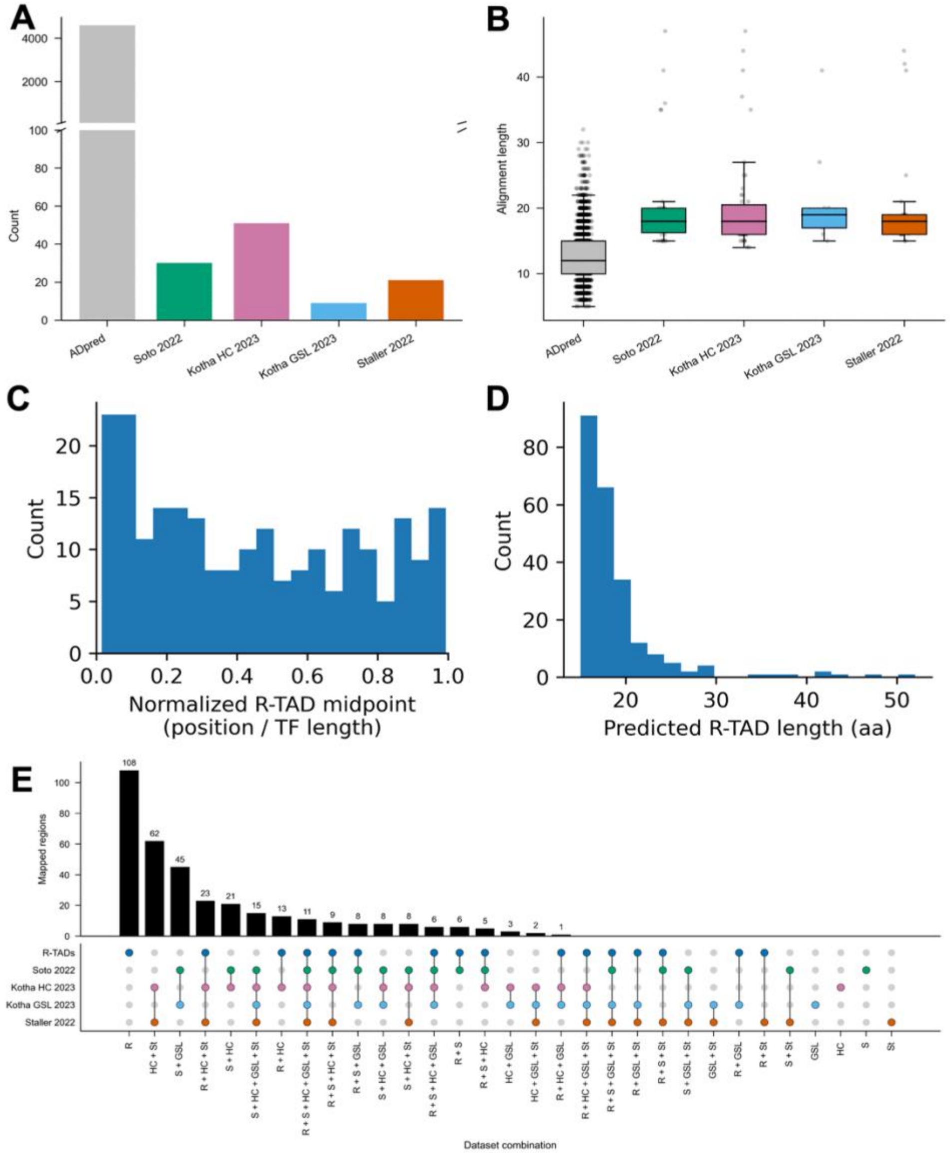


*Figure. S2 | Characterization of predicted R-TADs and sequence-based overlap with published activation-domain catalogues.*

(A) BLAST mapping yield across literature datasets: number of high-confidence same-protein matches between predicted R-TAD fragments and each activation-domain catalogue. A broken y-axis is used, with the lower axis spanning 0–100, to display smaller catalogues alongside the larger ADpred set on a linear scale.
(B) Alignment-length distributions for significant BLAST matches in each dataset.
(C) Length distribution of predicted R-TAD segments in the 100-TF prediction set.
(D) Distribution of predicted R-TAD positions along TF sequences using normalized segment midpoints (midpoint/TF length).
(E) UpSet-style summary of sequence-based overlap between predicted R-TAD regions and published AD/TAD datasets. Left, total number of mapped regions per

dataset. Right, overlap classes across datasets, with bars indicating the number of mapped regions in each intersection and the dot matrix indicating dataset membership for that intersection.

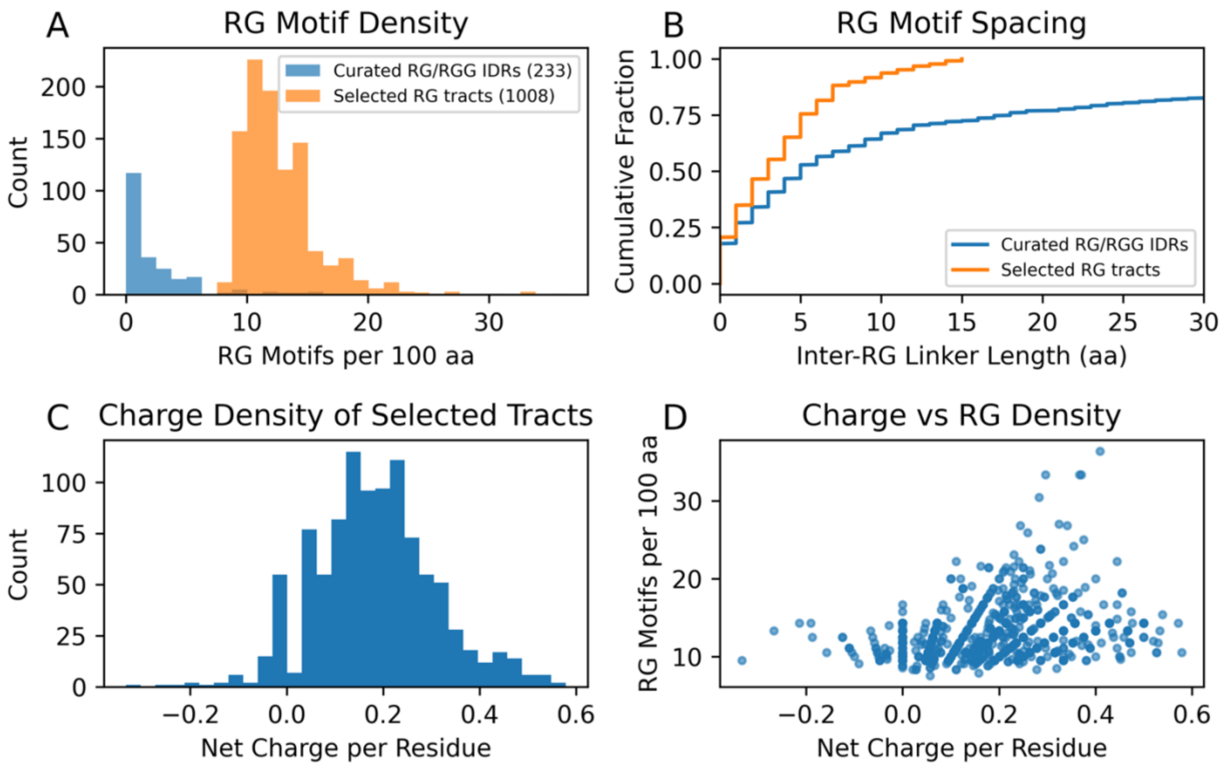


*Figure S3 | Motif density, spacing, and charge properties of compact RG regions used for TF–RBP pairing analyses.*

(A) Distribution of RG dipeptide density (RG motifs per 100 residues) comparing the 1,008 compact RG regions identified across the proteome to disordered regions derived from curated RG/RGG-containing proteins.

(B) Cumulative distributions of inter-RG linker lengths comparing compact RG regions to curated RG/RGG disordered regions, highlighting enrichment of short spacers in the compact-tract set.

(C) Distribution of net charge density (net charge per residue) across the 1,008 compact RG regions.

(D) Relationship between charge density (net charge per residue) and RG motif density (RG motifs per 100 residues) across compact RG regions.

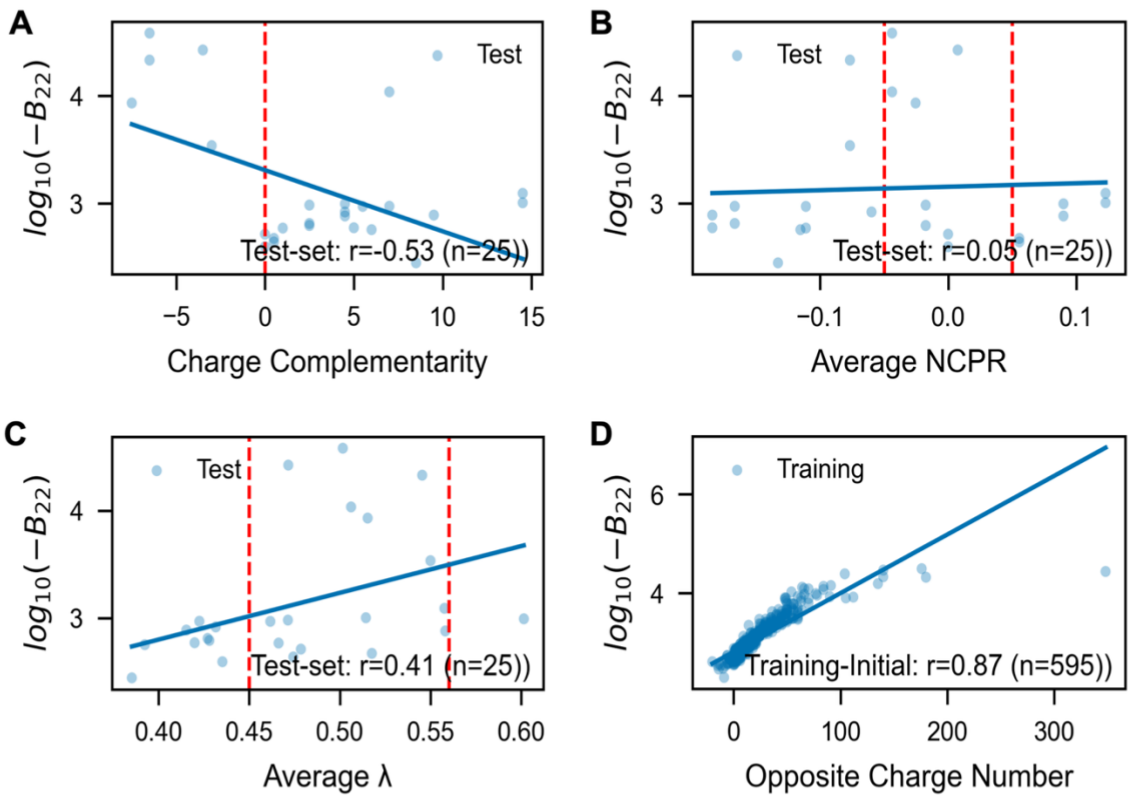


*Supplementary Fig. S4 | Motif density, spacing, and charge properties of compact RG regions used for TF–RBP pairing analyses.*

(A) Distribution of RG dipeptide density (RG motifs per 100 residues) comparing the 1,008 compact RG-rich regions identified across the proteome to disordered regions derived from curated RG/RGG-containing proteins.

(B) Cumulative distributions of inter-RG linker lengths comparing compact RG regions to curated RG/RGG disordered regions, highlighting enrichment of short spacers in the compact-tract set.

(C) Distribution of net charge density (net charge per residue) across the 1,008 compact RG regions.

(D) Relationship between charge density (net charge per residue) and RG motif density (RG motifs per 100 residues) across compact RG regions.

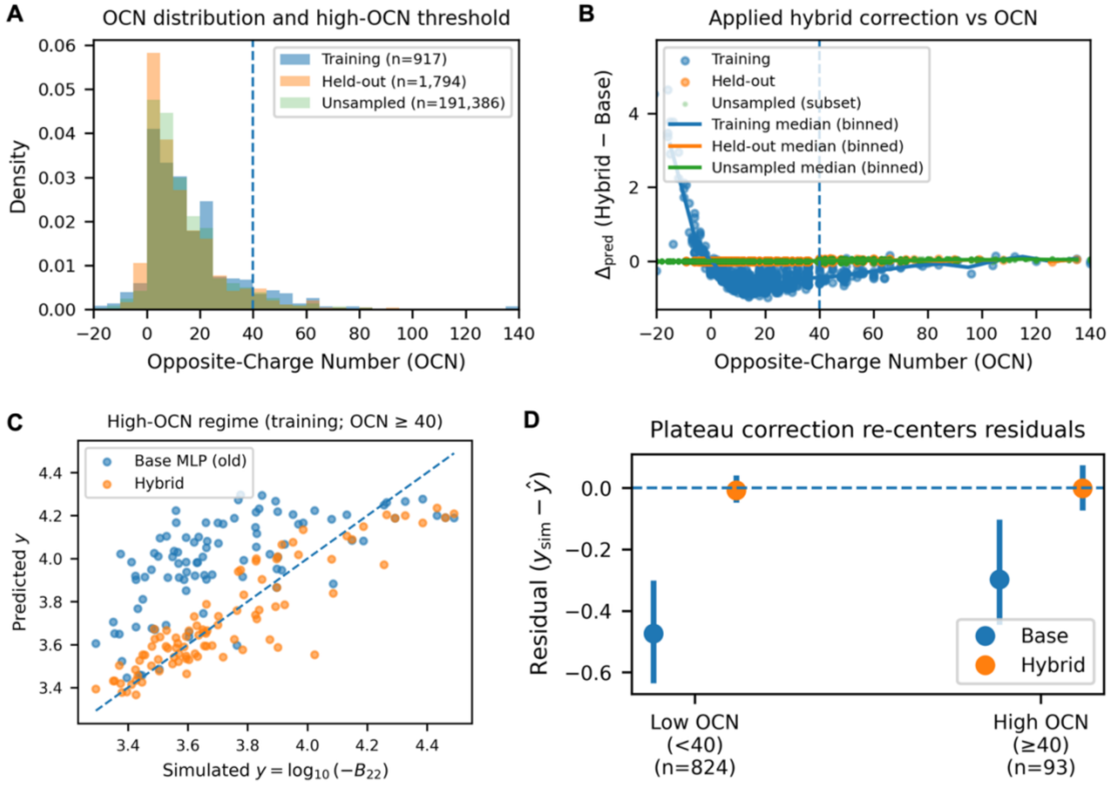


*Figure S5 | Characterization of the plateau-aware correction.*

(A) Distribution of opposite-charge number (OCN) across the training (n = 917), held-out (n = 1,794), and unsampled (n = 191,386) pair sets. The dashed vertical line marks the high-OCN threshold (OCN = 40) used to train and gate the residual correction.

(B) Magnitude of the hybrid correction as a function of OCN, shown as $\Delta_{pred}$=$y_{hybrid}$-$y_{base}$for training, held-out, and a random subset of unsampled pairs (for visualization). Solid lines show binned medians; the dashed line indicates OCN = 40. The correction is minimal at low OCN and becomes appreciable primarily in the high-OCN regime, consistent with the plateau-aware design.

(C) High-OCN training subset (OCN ≥ 40) comparing base and hybrid predictions against simulated $y$=$log_{10}$(-$B_{22}$). The dashed diagonal indicates the identity line. The hybrid model expands the dynamic range in the plateau regime relative to the base predictor.

(D) Residual bias by OCN regime (training set). Residuals ($y_{\mathrm{sim}}$-$\hat{y}$)are summarized separately for low OCN (<40) and high OCN (≥40). Points show median residuals and whiskers indicate the interquartile range (25th–75th percentile). The dashed horizontal line at zero indicates unbiased predictions. The hybrid model shifts residuals toward zero, consistent with a plateau-targeted correction.

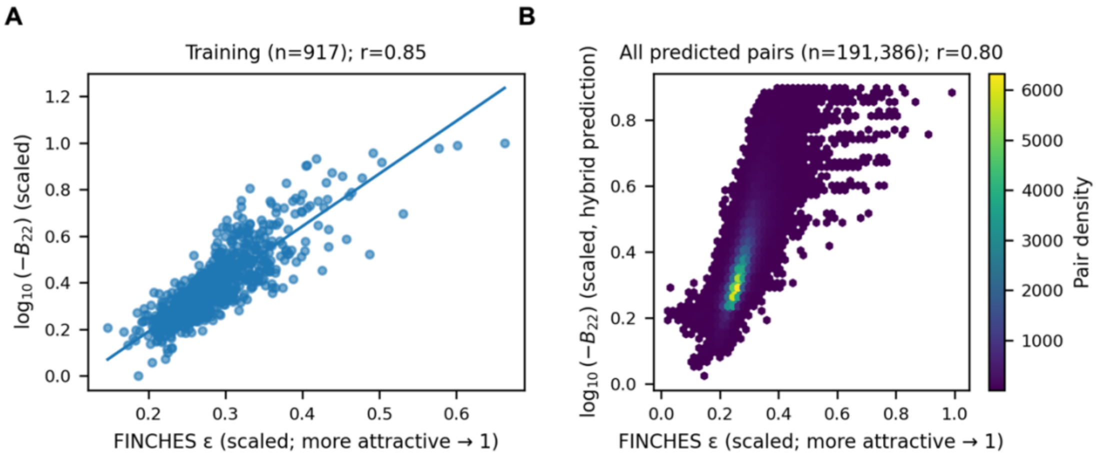


*Figure S6 | FINCHES interaction scores correlate with simulated and predicted interaction strength across the R-TAD–RGG space.*

(A) Training set (n = 917): scaled FINCHES $\varepsilon$ values plotted against scaled simulated interaction strength $y = log_{10}(-B_{22})$. FINCHES $\varepsilon$was transformed to an "attraction-scaled" variable in $[0,1]$so that more attractive (more negative) $\varepsilon$ corresponds to larger values: $\varepsilon_{scaled} = (\varepsilon_{max} - \varepsilon)/(\varepsilon_{max} - \varepsilon_{min})$. Interaction strength was min–max scaled to $[0,1]$ using global bounds shared across panels. Pearson correlation is reported in the panel title.

(B) All predicted pairs (n = 191,386): scaled FINCHES $\varepsilon$ plotted against scaled hybrid-predicted $y$, shown as a density hexbin. Color indicates pair density. Pearson correlation is reported in the panel title.

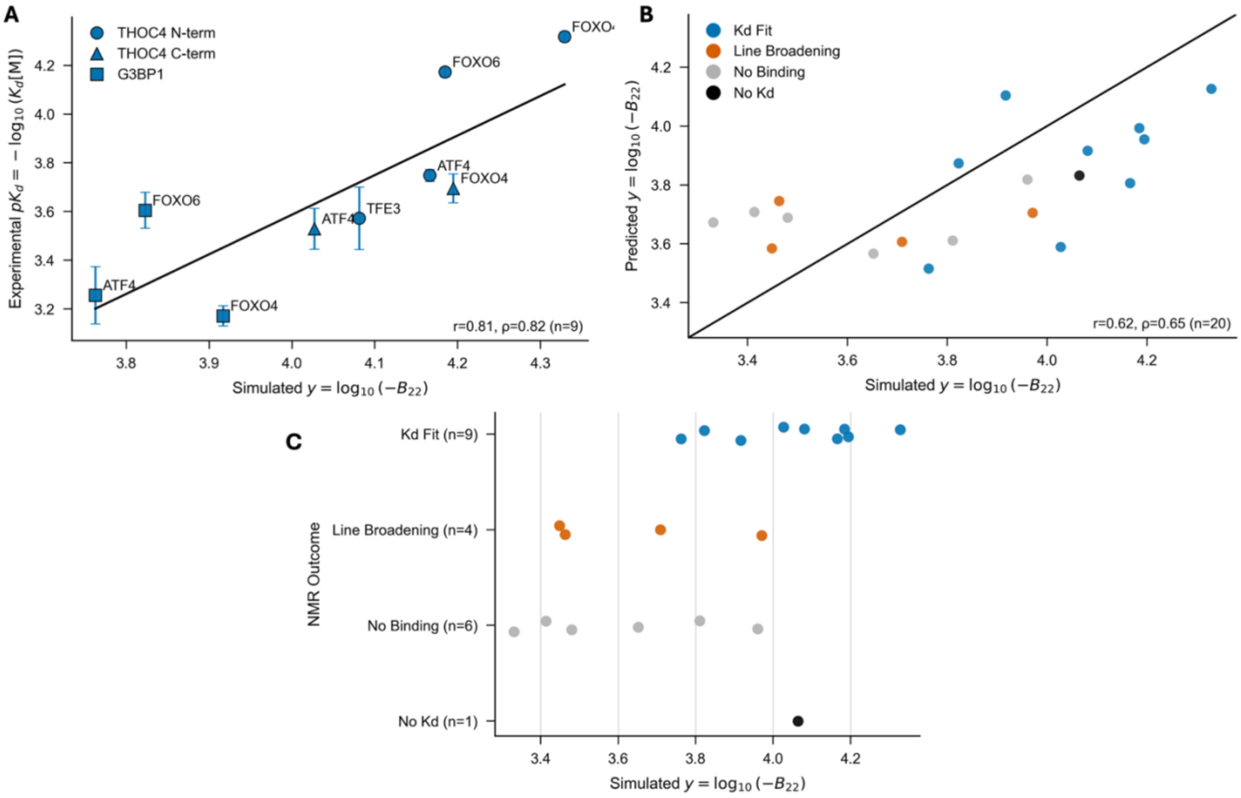


*Figure S7 | Simulated interaction strengths for the NMR validation set.*

(A) Simulated interaction strength $y{=}log_{10}(\text{-}B_{22})$versus experimental $pK_d$for Kd-fit interactions. $pK_d$values were computed from TITAN-derived Kd (µM) as $pK_d{=}6\text{-}log_{10}(K_d[\mu M])$; error bars reflect propagated TITAN fit errors.

(B) Predicted $y{=}log_{10}(\text{-}B_{22})$versus simulated $y{=}log_{10}(\text{-}B_{22})$for all tested pairs, coloured by experimental outcome class.

(C) Distribution of simulated interaction strengths across outcome categories (Kd fit, line broadening, no binding), showing separation of outcome classes along the simulated interaction scale.
